# Bacterial cell cycle control by citrate synthase independent of enzymatic activity

**DOI:** 10.1101/799742

**Authors:** Matthieu Bergé, Julian Pezzatti, Víctor González-Ruiz, Laurence Degeorges, Serge Rudaz, Patrick H. Viollier

## Abstract

Coordination of cell cycle progression with central metabolism is fundamental to all cell types and likely underlies differentiation into dispersal cells in bacteria. How central metabolism is monitored to regulate cell cycle functions is poorly understood. A forward genetic selection for cell cycle regulators in the polarized alpha-proteobacterium *Caulobacter crescentus* unearthed the uncharacterized CitA citrate synthase, a TCA (tricarboxylic acid) cycle enzyme, as unprecedented checkpoint regulator of the G1→S transition. We show that loss of the CitA protein provokes a (p)ppGpp alarmone-dependent G1-phase arrest without apparent metabolic or energy insufficiency. While S-phase entry is still conferred when CitA is rendered catalytically inactive, the paralogous CitB citrate synthase has no overt role other than sustaining TCA cycle activity when CitA is absent. With eukaryotic citrate synthase paralogs known to fulfill regulatory functions, our work extends the moonlighting paradigm to citrate synthase coordinating central (TCA) metabolism with development and perhaps antibiotic tolerance in bacteria.

## INTRODUCTION

Nutritional control on cellular development and cell cycle progression have been described in many systems, but only in few instances are the molecular determinants known that govern the responses. Bacteria are attractive models to elucidate the underlying mechanism because of their genetic tractability, their apparent morphological and cellular simplicity, and the robust influence of the changing nutritional states on their growth and morphology. Several links between central metabolism and cell-cycle have been described (Monahan and Harry 2016) but the underlying molecular mechanisms are poorly understood. Several metabolic enzymes, often enzyme paralogs, are known to be appropriated for regulatory functions, in place or in addition to their normal enzymatic functions. These proteins, called moonlighting or trigger enzymes, are ideal coupling factors to coordinate regulatory changes in response to metabolic fluctuations (Huberts and van der Klei 2010; Commichau and Stülke 2015). A notorious example is the aconitase paralog IRE-BP that fulfills a regulatory function as mRNA repressing protein.

The synchronisable α-proteobacterium *Caulobacter crescentus* is the preeminent model to elucidate cell cycle control mechanisms. Cell division in *C. crescentus* is asymmetric and thus yields two dissimilar daughter cells: a stalked and capsulated S-phase cell that replicates its genome before dividing, and a piliated and flagellated dispersal (swarmer) cell that resides in the non-replicative and non-dividing G1-phase (Figure 1A) (Goley et al. 2007; Kirkpatrick and Viollier 2012; Ausmees and Jacobs-Wagner 2003). While the old pole of the stalked cell features a cylindrical extension of the cell envelope (the stalk), the one of the swarmer cell is characterized by a flagellum and several adhesive pili. The polar placement of organelles is dictated by polar scaffolding proteins including the TipN coiled-coil protein (Figure 1A) (Huitema et al. 2006; Lam et al. 2006) and the PopZ polar organizer (Bowman et al. 2008; Ebersbach et al. 2008). As polar remodeling occurs during the cell cycle, it is not surprising that polarity determinants also affect cell cycle progression (Bergé and Viollier. 2017).

**Figure 1.**
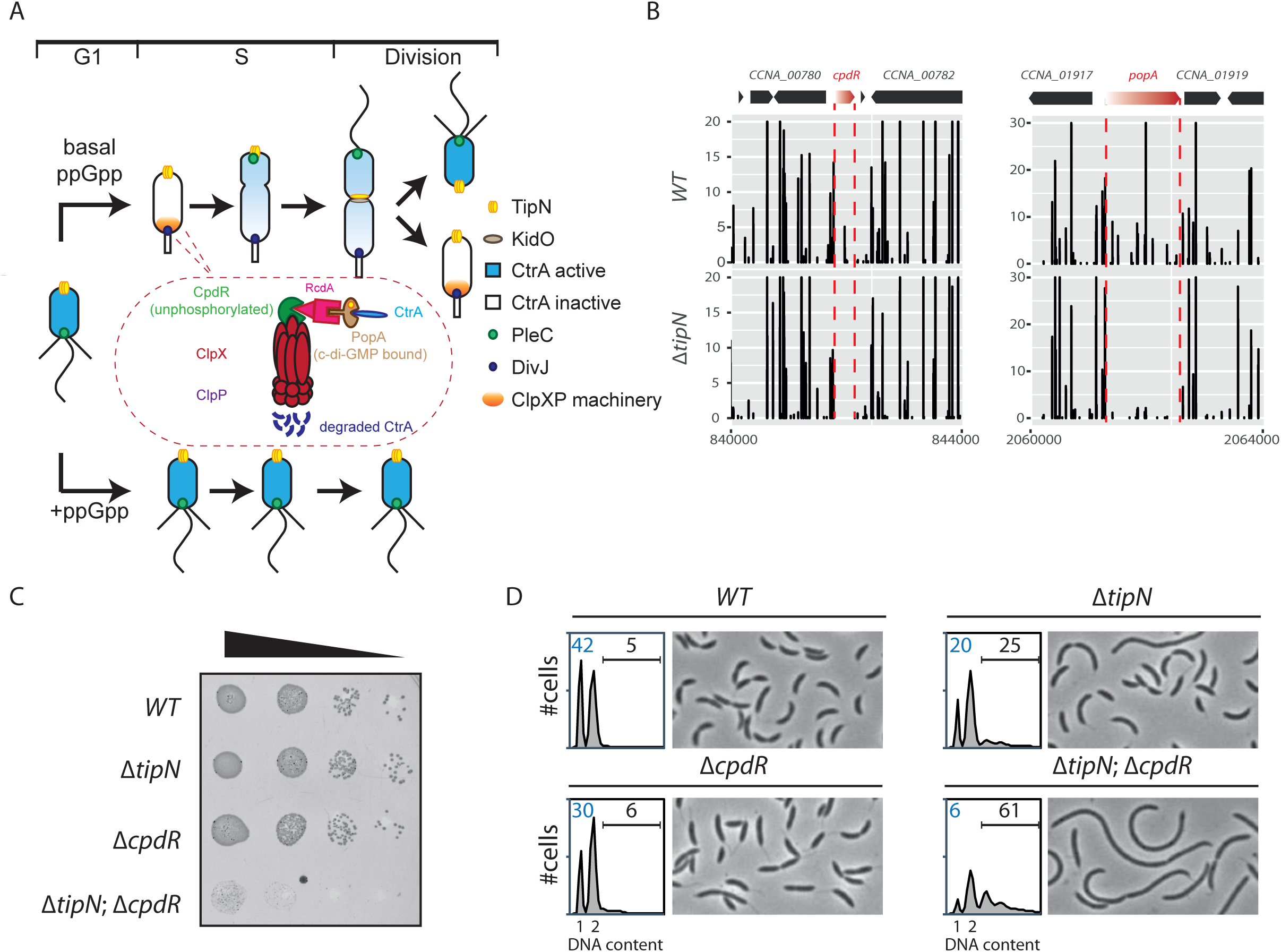
Synthetic sick interaction between *tipN* and the adaptors of the ClpXP machinery. (A) Schematic of the different stage of *C. crescentus* cell cycle (G1 phase, S phase and division are showed) in normal condition (upper part). TipN (yellow dot) and KidO (brown circle) localization are represented along the cell cycle. Phosphorylated CtrA (blue) activates transcription of G1 phase genes and prevent DNA replication in the swarmer cell. Upon transition from a swarmer to stalked cell, ClpXP machinery (red) and its adaptor CpdR (green), RcdA (pink) and PopA (beige) localizes to the incipient stalked pole where it degrades CtrA, allowing DNA replication and cell division. In the pre-divisional cell, the antagonistic kinase/phosphatase pair, DivJ (purple dot) and PleC (green dot) indirectly influence the phosphorylation of CtrA with the stalked cell compartment or swarmer cell compartment respectively. PleC promotes CtrA phosphorylation in the swarmer cell while DivJ prevents its phosphorylation in the stalked cell. Pili and flagella are depicted as straight wavy lines respectively. (p)ppGpp production occurring in carbon or nitrogen starvation prevents swarmer to stalked cell transitions (bottom part). (B) Transposon libraries were generated in the *WT* and the Δ*tipN* mutant (MB556). The sites of Tn insertion were identified by deep sequencing and mapped onto the *C. crescentus* NA1000 reference genome. Two regions of the genome showing the *cpdR* and *popA* locus are depicted. The height of each line reflects the number of sequencing reads at this position and all the graph between *WT* and Δ*tipN* are scaled similarly. Tn insertions in *cpdR* and *popA* were reduced in the Δ*tipN* mutant compared to the *WT*. (C) Spot dilutions of the indicated strains (MB1, MB556, MB2001, MB2017 from top to bottom). The four strains were grown overnight, adjusted at an OD_600_ of 0.5 and serially diluted. Eight microliters of each dilutions were spotted onto PYE plates. (D) Flow cytometry profiles and phase contrast images of *WT* (MB1), Δ *tipN* (MB556), Δ*cpdR* (MB2001) or Δ*tipN* Δ*cpdR* (MB2017) double mutants. Genome content (labelled as DNA content) was analyzed by FACS during exponential phase in PYE.

The swarmer cell is obliged to differentiate into a stalked cell in order to generate progeny. During the swarmer-to-stalked cell transition (also known as the G1→S transition), the flagellum is shed, pili are retracted, and a stalk is elaborated from the vacated pole while replication competence is acquired. A regulatory protein that coordinates morphological and cell cycle stages is the essential <u>c</u>ell cycle transcriptional <u>r</u>egulator <u>A</u>, CtrA, a DNA-binding (OmpR-like) response regulator that upon phosphorylation, directly binds and regulates the origin of replication (*ori*) (Laub et al. 2000; Quon et al. 1996) and promoters of developmental genes, including promoters that fire only in G1-phase (Fumeaux et al. 2014), such as *pilA* which encodes the structural subunit of the pilus filament (Skerker and Shapiro 2000), several flagellin genes and other genes controlling cell envelope modification (Ardissone and Viollier 2015).

CtrA is phosphorylated (CtrA∼P) by a complex phospho-signaling pathway that regulates the activity and abundance of CtrA during the *C. crescentus* cell cycle (Figure 1A) (Jacobs et al. 1999; Biondi et al. 2006; Paul et al. 2008; Wu et al. 1998; Tsokos and Laub 2012). During the G1→S transition, CtrA is dephosphorylated and a proteolytic branch responsible for the degradation of CtrA is activated (Joshi and Chien 2016). This pathway involves the protease ClpXP and three selectivity factors that present CtrA to ClpXP (Figure 1A). These proteolytic adaptors namely CpdR, RcdA and PopA are organized into a regulatory hierarchy that coordinates the degradation of many substrates precisely during the G1→S transition (Iniesta and Shapiro 2008; McGrath et al. 2006; Duerig et al. 2009; Joshi et al. 2015). Upon degradation of CtrA, the DNA replication block is relieved and G1-phase genes are no longer expressed. Thus, maintenance of cells in G1 phase, requires that CtrA remains present and phosphorylated (Domian et al. 1997).

The duration of the G1 period is affected by nutrient availability in *C. crescentus* and other α-proteobacteria (De Nisco et al. 2014). Upon nitrogen or carbon starvation, the G1→S transition is blocked (Leslie et al. 2015; England et al. 2010; Lesley and Shapiro 2008; Britos et al. 2011; Gorbatyuk and Marczynski 2005). This G1 block is associated with the accumulation of the guanosine tetra-and penta-phosphate [(p)ppGpp] alarmone (Figure 1A) (Lesley and Shapiro 2008; Ronneau et al. 2016), which affects important cellular processes in bacteria such as transcription, translation or DNA replication (Liu et al. 2015; Zhang et al. 2018; Wang et al. 2019). Rsh family proteins directly modulate the intracellular level of (p)ppGpp and most bacterial genomes encode at least one bifunctional Rsh protein able to synthesize and hydrolyze (p)ppGpp. *C. crescentus* encodes a single bifunctional enzyme, named SpoT (Lesley and Shapiro 2008; Ronneau et al. 2016; Atkinson et al. 2011; Boutte et al. 2012). Previous studies have shown that (p)ppGpp accumulation leads to a stabilization of CtrA by an unknown, yet (p)ppGpp-dependent mechanism, impairing the G1→S transition (Leslie et al. 2015; Lesley and Shapiro 2008; Gonzalez and Collier 2014).

Here, we report that citrate synthase (CitA), the first enzyme of the Krebs (tricarboxylic acid, TCA) cycle that condenses oxaloacetate and acetyl-CoA, fulfills an unprecedent role as a checkpoint regulator controlling the G1→S transition. Selecting for mutations that elevate the G1-phase population unearthed a loss of function-mutation in the *citA* gene. The effects of this mutation are nullified when (p)ppGpp production is lost and are not due to glutamate auxotrophy, as it is typically the case for citrate synthase mutants in other bacterial model systems such as *Escherichia coli*. Even though CitA is a functional citrate synthase, the paralog CitB can sustain enzymatic function but not cell cycle control. As even catalytically inactive CitA retains cell cycle control, we conclude that it acts as the first bacterial moonlighting enzyme that acts on central metabolism and, independently, on S-phase entry.

## RESULTS

### G1-phase defect in cells lacking TipN and adaptors of the ClpXP machinery

Flow cytometric analysis by fluorescence activated cell sorting (FACS) is a convenient way of scoring cell cycle defects of a population of cells. In our efforts to explore the function of the TipN polarity factor, we conducted FACS analysis of wild-type (*WT*) and *ΔtipN* cells and found a 47% reduction in the abundance of G1-phase cells (Figure 1D) in the latter, without a strong effect on growth or efficiency of plating (Figure 1C). Next, we sought mutations that accentuate the G1-phase defect of *ΔtipN* by comparative transposon (Tn) insertion site sequencing (Tn-Seq) of *WT* and *ΔtipN* cells, reasoning that Tn insertions that decrease the fitness of *ΔtipN* cells might further reduce the production of G1-phase cells. This Tn-Seq analysis revealed that Tn insertions in the *popZ* gene encoding a polar scaffold protein (Bowman et al. 2008; Ebersbach et al. 2008) were underrepresented in Δ*tipN* compared to *WT* cells (Supplemental Table 1), recapitulating the synthetic lethal interaction between the two polarity hubs encoded by *tipN* and *popZ* (Ebersbach et al. 2008). Surprisingly, Tn-Seq also revealed that Tn insertions in genes reducing CtrA activity (PleC) are underrepresented in Δ*tipN versus WT* cells while genes increasing CtrA activity are overrepresented (DivJ), suggesting that the activity of the G1-phase regulator CtrA is reduced in Δ*tipN* cells as already hinted by FACS analysis (see above).

Paradoxically, our Tn-Seq analysis also revealed an underrepresentation of Tn insertions in the *cpdR*, *rcdA* and *popA* genes in *ΔtipN versus WT* cells (Figure 1B and Figure1-Figure supplemental 1A), a result that was confirmed by reverse Tn-Seq experiment determining the abundance of Tn insertions in the *tipN* gene of Δ*cpdR versus WT* cells (Figure 1- Figure supplemental 1B). If CtrA activity is low in Δ*tipN* cells, then Tn insertions in *cpdR*, *rcdA* or *popA* would be expected to have a beneficial effect to Δ*tipN* cells because these genes promote the turnover of CtrA (and other proteins) at the G1àS transition (Joshi and Chien 2016; Joshi et al. 2015; Iniesta et al. 2006; Duerig et al. 2009; McGrath et al. 2006), so disruption in these genes should raise CtrA levels. Alternatively, since they also control the stability of proteins other than CtrA, another ClpXP substrate might be responsible for enhancing the cell cycle defect of Δ*tipN* cells.

To confirm the genetic relationship between *tipN* and *cpdR, rcdA* or *popA*, we created double mutants by introducing either the Δ*cpdR*, Δ*rcdA* or Δ*popA* mutation into Δ*tipN* cells and found that all double mutants exhibit a reduction in viability by three orders of magnitude on a logarithmic scale (Figure 1C; Figure 1- Figure supplemental 1C and 1D). Examination of Δ*tipN* Δ*cpdR* double mutant cells by phase contrast microscopy revealed that they are 70% more elongated on average compared to *WT* and Δ*cpdR* or Δ*tipN* single mutant cells (Figure 1D and Figure 1- Figure Supplemental 1F). Flow cytometry analysis of exponentially growing Δ*tipN* Δ*cpdR* double mutant cells by fluorescence activated cell sorting (FACS) revealed a strong reduction (85%) in the number of G1-phase cells and a massive increase of cells with multiple (>2) chromosomes compared to *WT* cells, whereas Δ*cpdR* and Δ*tipN* single mutants only showed a slight decrease in the G1 population (Figure 1D). Importantly, the Δ*tipN* Δ*rcdA* and Δ*tipN* Δ*popA* double mutants showed a similar accumulation of elongated cells and reduction in G1-phase cells number (Figure 1- Figure supplemental 1E and 1F).

We reasoned that the accumulation of a ClpXP substrate whose degradation is dependent on CpdR, RcdA and PopA causes a cell cycle defect in Δ*tipN* cells. Seeking to uncover Tn insertions in a gene encoding a ClpXP substrate that when inactivated ameliorates growth of Δ*tipN* Δ*cpdR* cells, we conducted Tn-Seq in Δ*tipN* Δ*cpdR* double mutant cells and found a 19-fold increase in Tn insertions in the *kidO* gene in Δ*tipN* Δ*cpdR* double mutant cells compared to *WT* cells or Δ*tipN* and Δ*cpdR* single mutant cells (Figure 1- Figure supplemental 2A). KidO is a bifunctional oxidoreductase-like protein degraded by ClpXP that integrates cell fate signaling with cytokinesis in *C. crescentus* (Radhakrishnan et al. 2010) by preventing premature assembly of the cytokinetic structure of FtsZ polymers in G1 phase and promoting FtsZ-ring disassembly in G2 phase (Beaufay et al. 2015). KidO is not present in S-phase when FtsZ polymerizes at the division site as its degradation by ClpXP is stimulated by CpdR, RcdA and PopA at the G1→S transition. In cells lacking these proteolytic adaptors, KidO is no longer degraded at the G1→S transition and is, therefore, present throughout the cell cycle (Radhakrishnan et al. 2010). To test if KidO stabilization induces filamentation of Δ*tipN* Δ*cpdR* cells, we expressed the *kidO^AA::DD^* allele from the *xylX* locus in Δ*tipN* cells. This allele encodes a mutant form of KidO in which the two penultimate alanine residues are both substituted by aspartic acid, a double substitution that prevents degradation of KidO by the ClpXP protease at the G1→S transition, akin to the Δ*cpdR* mutation (Radhakrishnan et al. 2010). The resulting Δ*tipN xylX*::*kidO^AA::DD^* cells are highly filamentous, even without induction of the *xylX* promoter by xylose, with a strong decrease of the G1 population and an increase of cells containing more than two chromosomes recapitulating the phenotype of the Δ*tipN* Δ*cpdR* double mutant cells (Figure 1- Figure supplemental 2B). Conversely, an in-frame deletion in *kidO* (Δ*kidO*) restores a near *WT* division phenotype to Δ*tipN* Δ*cpdR* cells strain (Figure 1- Figure supplemental 2B).

Thus, stabilization of KidO in cells lacking TipN inhibits cell division and prevents the accumulation of G1 dispersal cells.

### Genetic screen to identify regulators of the G1 to S transition

Since KidO is also known to act negatively on CtrA phosphorylation which is required for G1 cell accumulation (Radhakrishnan et al. 2010), we speculated that decrease in the G1 population of Δ*tipN* Δ*cpdR* is due to very low CtrA activity. To confirm that CtrA activity is indeed reduced, we introduced a *pilA::*P*pilA*-*GFP* probe reporter into the *pilA* locus of *WT*, *ΔtipN* or *ΔcpdR* single mutant, and *ΔtipN ΔcpdR* double mutant cells. This reporter harbors the CtrA-dependent *pilA* promoter (P*_pilA_*) that fires in G1-phase and PilA start codon translationally fused to the green fluorescent protein (GFP). GFP expression from this reporter can be conveniently observed and quantified by live-cell fluorescence microscopy (Figure 2A). In agreement with the FACS analysis shown in Figure 1E, GFP fluorescence is reduced in *ΔtipN* cells versus WT or *ΔcpdR* cells, but in *ΔtipN ΔcpdR* double mutant cells, a strong decrease in GFP fluorescence was observed indicating a strong downregulation in CtrA-dependent reporter activity. Likewise, a transcriptional fusion of P*_pilA_* to the promoter-less *nptII* gene (conferring resistance to kanamycin) at the *pilA* locus (*pilA*::P*_pilA_*-*nptII*) is strongly reduced in *ΔtipN ΔcpdR* double mutant cells *versus WT* cells, precluding growth of *ΔtipN ΔcpdR* on plates containing 20 µg/mL kanamycin (Figure 2B) because P*_pilA_* is poorly active, whereas *WT* cells harboring this reporter grow.

**Figure 2.**
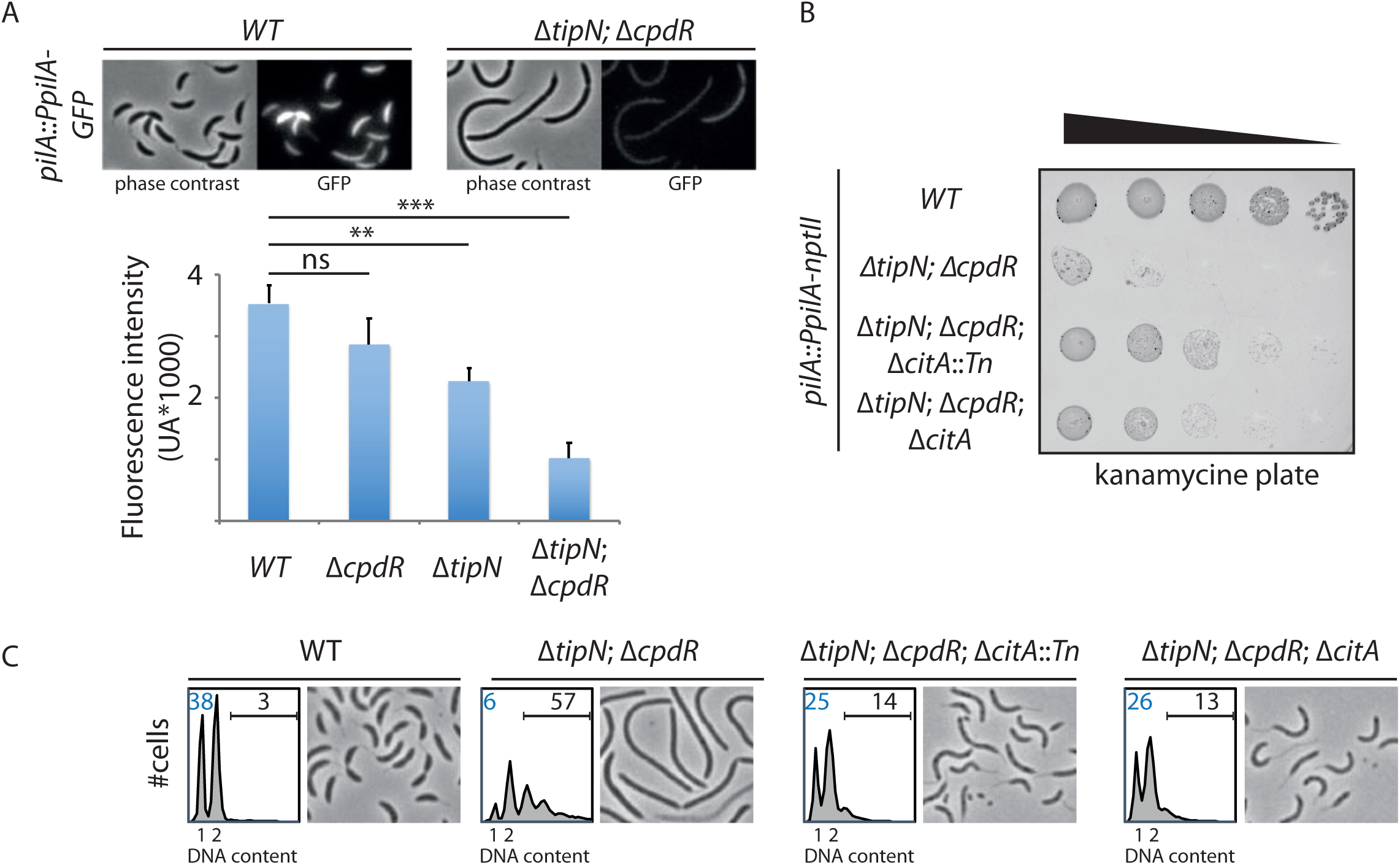
Genetic screen to identify Tn insertions that enhance CtrA. (A) CtrA activity in the *WT* (MB2325), Δ*tipN* (MB2337) and Δ*cpdR* (MB2329) single mutant cells, and Δ*tipN* Δ*cpdR* (MB2331) double mutant cells was monitored using a *pilA::P_pilA_-GFP* transcriptional reporter whose activity is dependent on the activity of CtrA. Fluorescence intensity was automatically quantified and t-test was performed to determine the significance with p<0.05 (**) and p<0.005 (***). (B) Spot dilutions of the indicated strains (MB2268, MB2271, MB3056, MB3058 from top to bottom) carrying the *pilA::P_pilA_-nptII* transcriptional reporter on PYE plates containing kanamycin (20µg.mL^-1^). (C) FACS profiles and phase contrast images of the strain described in panel B. FACS analysis showing genome content (ploidy) of cells growing exponentially in PYE and then treated by rifampicin (50 μg/ml) for 3 h to inhibit DNA replication. Numbers (%) of G1-phase cells and cells containing more than 2 chromosomes is indicated in blue and black respectively.

Next we used these reporter cells to find mutations that maintain CtrA active in the absence of TipN and CpdR. To this end, we mutagenized *ΔtipN ΔcpdR* P*_pilA_*-*nptII* reporter cells using a mini-*himar1* Tn (Mar2xT7), encoding gentamycin resistance (GmR), and selected for growth on plates containing kanamycin and gentamycin. Among several isolated mutants, one mutant was isolated harboring a Tn insertion in the middle of the *CCNA_01983* (henceforth *citA*) gene whose gene product is annotated as a type II citrate synthase (PRK05614). After backcrossing experiments confirmed that the *citA*::Tn mutation confers kanamycin resistance to *ΔtipN ΔcpdR* P*_pilA_*-*nptII* reporter cells, we engineered an in-frame deletion of *citA* (Δ*citA*) and found that this mutation also supports growth *ΔtipN ΔcpdR* P*_pilA_*-*nptII* reporter cells on kanamycin plates, indicating that loss of *citA* function augments P*_pilA_* activity (Figure 2B). The *citA*::Tn or the Δ*citA* mutations both correct the abnormal cell size distribution (cell filamentation) and augment the G1 population of *ΔtipN ΔcpdR* double mutant cells (Figure 2C and Figure 2- Figure supplemental 1).

In sum, inactivation of *citA* gene leads to an increase of P*_pilA_* activity and G1 cell production, while ameliorating the division defect of cells lacking TipN and CpdR.

### CitA encodes a citrate synthase

The primary structure of CitA resembles citrate synthases that execute the first enzymatic reaction in the Krebs (tricarboxylic, TCA) cycle with the condensation of the acetyl group from acetyl-CoA onto oxaloacetate to form citrate (Figure3-Figure supplemental 1A) (Figure 3A). *C. crescentus* CitA has 65% amino acid identity to the GltA citrate synthase from *Escherichia coli* K12 (strain MG1655) and 32 % identity to CitA from *Bacillus subtilis* (strain 168). To confirm that *C. crescentus* CitA indeed has citrate synthase activity, we probed for heterologous complementation of the glutamate auxotrophy of *E. coli* Δ*gltA* cells that lack citrate synthase activity (Lakshmi and Helling 1976). To this end, we engineered *E. coli ΔgltA* cells expressing either *C. crescentus* CitA or *E. coli* GltA from a multicopy plasmid. As expected, *E. coli ΔgltA* cells harboring the empty vector are unable to grow in (M9) minimal medium without glutamate, but *ΔgltA* cells grew well in the presence of either the *gltA-* or the *citA*-expression plasmid (Figure 3B). Thus*, C. crescentus citA* encodes a functional citrate synthase.

**Figure 3.**
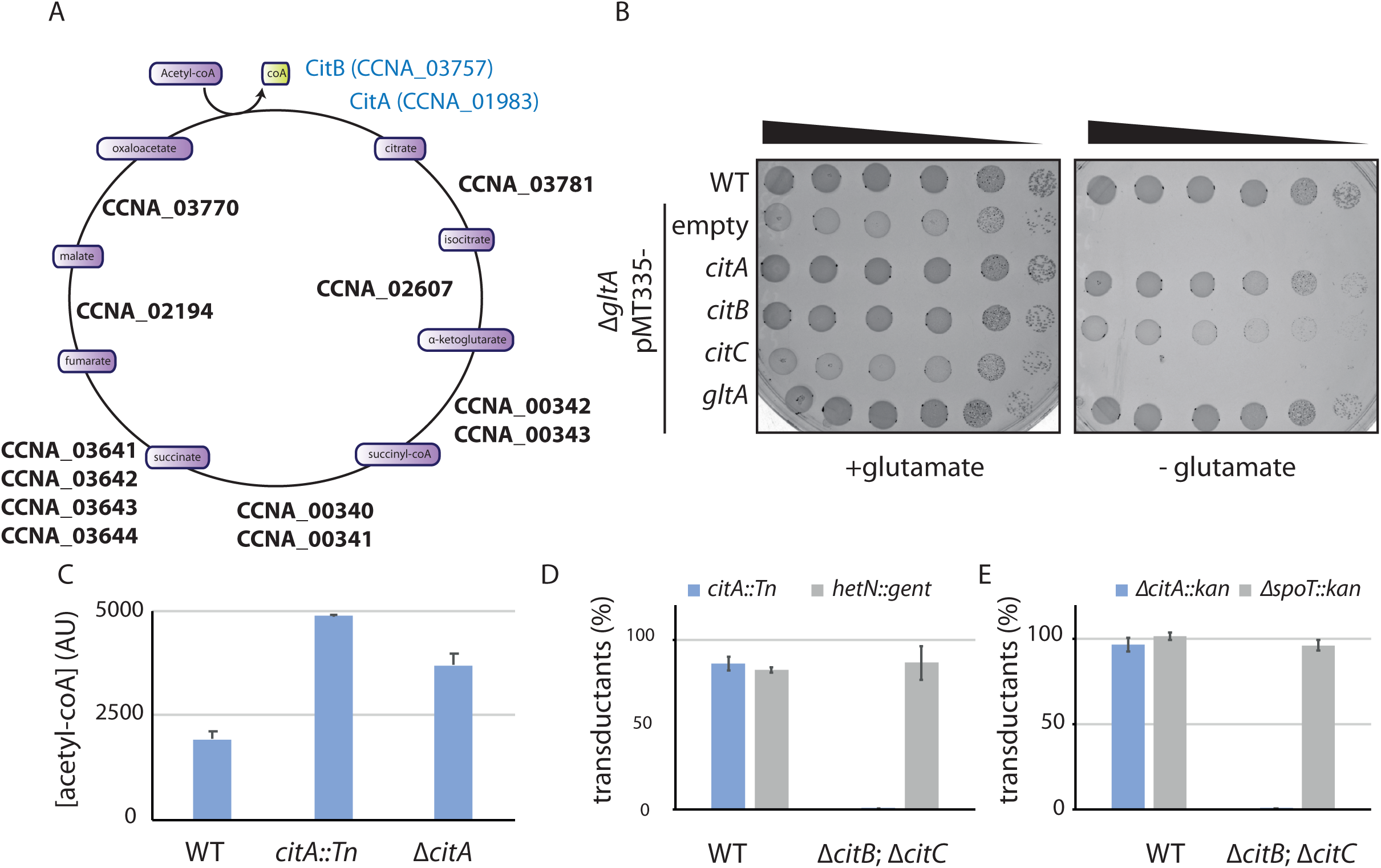
*C. crescentus* genome encodes two functional citrate synthase. (A) A schematic Krebs cycle is represented. The two functional citrate synthase are indicated in blue. Essential enzymes from the Krebs are highlighted in bold. (B) Spot dilutions of the indicated *E. coli* strains (eMB554, eMB556, eMB558, eMB560, eMB562 and eMB564 from top to bottom) on minimal medium containing or not glutamate. Only the strain carrying a functional citrate synthase can grow without glutamate. (C) LC-MS-based quantification of acetyl-CoA in extract of *WT* (MB1), *citA*::Tn (MB2622) and Δ*citA* (MB2559) cells grown in PYE liquid cultures. Error bars denote the standard deviation from three biological replicates. (D) ϕCr30-mediated generalized transduction frequencies of *citA*::Tn into WT (MB1) or Δ*citBC* double mutant cells (MB2679). For transduction, cells were normalized according to the OD_600_ ∼1and infected with the same amount of ϕCr30 harboring either *citA*::Tn or a transposon insertion in the *hetN* gene (encoding gentamycin resistance) as a control of transduction-The transductions were selected on PYE plates containing gentamycin. The numbers of colonies were counted after 3 days of incubation at 30°C. Error bars denote the standard deviation for three independent experiments. Cells harboring the Δ*citBC* mutation are not able to accept *citA*::Tn mutation. (E) Same as in panel D using the Δ*citA::kan* allele or a deletion in the *spoT* gene (encoding kanamycin resistance) delivered by ϕCr30-mediated generalized transduction as a control. Transductants were selected on PYE plates containing kanamycin.

Next, we conducted metabolic profiling experiments using liquid chromatography coupled to high-resolution mass spectrometry (LC-HRMS) to quantify the abundance of intracellular metabolites in *C. crescentus WT* and *citA*::Tn or Δ*citA* cells grown in PYE (Pezzatti et al. 2019a). Robust quantitation of 103 metabolites (Supplemental Table S2) revealed that the metabolomic profile of *citA*::Tn resembles that of *ΔcitA* cells. Surprisingly, these metabolomic analyses did not show any significant difference in many of TCAs like citrate and isocitrate (Figure 3- Figure supplemental 1B). An indication that TCA cycle flux is nevertheless affected in the absence of CitA comes from the observation of a small increase in the levels of acetyl-CoA, as would be expected for citrate synthase mutant cells (Figure 3C).

The relatively modest effect of the *ΔcitA* mutation on TCA cycle activity might be due to the presence of a protein(s) other than CitA with citrate synthase activity. Unlike other TCA cycle enzymes, CitA is not essential for viability of *C. crescentus* cells on PYE (Christen et al. 2011). Therefore, we reasoned that CitA is not the only citrate synthase-like protein encoded in the *C. crescentus* genome. Indeed, BLAST searches revealed the presence of two other putative citrate synthase genes: *CCNA_03757* and *CCNA_03758* (Figure 3- Figure supplemental 1A) (henceforth *citB* and *citC*, respectively), annotated also as non-essential for viability (Christen et al. 2011). The *citB* and *citC* genes encode proteins with 30% and 32% identity to CitA from *C. crescentus*, 30% and 33% identity to GltA from *E. coli* K12 (MG1655) and 37% and 32% identity to CitA from *B. subtilis* (168). We therefore tested the ability of *citB* and *citC* to support citrate synthase function by heterologous complementation of the glutamate auxotrophy of *E. coli ΔgltA* cells on minimal medium lacking glutamate and found that expression of CitB, but not CitC, supported growth (Figure 3B). Thus, *C. crescentus citB* also encodes a functional citrate synthase. On the basis of these results, we speculated that *C. crescentus citA* mutants are able to grow because of residual citrate synthase activity conferred by CitB. To test if CitA is essential in cells lacking both *citB* and *citC*, we first created a strain with in-frame deletions in *citB* and *citC* (Δ*citBC*) and then attempted to introduce *citA*::Tn (encodes gentamycin resistance) or *ΔcitA* (tagged with a kanamycin resistance marker, *ΔcitA*::pNPTS138) by fCr30-mediated generalized transduction. Unlike *WT* cells, Δ*citBC* cells did not accept *citA*::Tn or *ΔcitA*::pNPTS138 generalized transducing particles (Figure 3D), but accepted generalized transducing particles harboring another genomic locus marked with either the gentamycin or the kanamycin resistance gene with similar efficiency as *WT* cells. We conclude that *C. crescentus* encodes at least two functional citrate synthases, one of which is absolutely required for growth on PYE.

### CitA promotes S-phase entry

To determine how loss of CitA signals G1 cell accumulation, we combined population-based and single cell approaches. First, EOP and growth curves indicate that the absence of CitA leads to a slow growth phenotype on PYE rich medium and that CitA is required for growth on minimal M_2_G medium (Figure 4A). Phase contrast microscopy of *citA*::Tn or Δ*citA* mutants showed that they are more phase-bright than *WT* cells (Figure 4B). In *C. crescentus* phase darkness is typically caused by intracellular polyphosphate granules that appear under stress conditions (Boutte et al. 2012). Thus, *citA* mutant cells might be defective in accumulating polyphosphate granules, perhaps because they are metabolized [converted into (p)ppGpp, see below] when cells are under nutritional stress. Moreover, microscopy revealed that Δ*citA* cells are shorter and narrower than *WT* cells (area of 0.42±0.009 µm and 0.43±0.007 µm respectively for the *citA*::Tn and Δ*citA* compared to 0.69±0.01 µm for *WT* cells, Figure 4B), perhaps because they spend more time in the non-growing G1 phase. Indeed, FACS analysis revealed a strong increase in the G1-phase population in the absence of CitA (68.3±1.25% and 69.3±1.22 of *citA*::Tn and Δ*citA* cells reside in G1 phase compared to 36.1±0.6% of *WT* cells, Figure 4B). Importantly, these phenotypes of *citA* mutant cells cannot be corrected by the addition of exogenous glutamine and, therefore, are not related to glutamine auxotrophy. Indeed, addition of glutamine to PYE or to M2G (minimal medium) did not ameliorate growth or division as determined by EOP assays (Figure 4- Figure supplemental 1A). Moreover, the addition of glutamine did also not restore a normal FACS profile to *citA* mutant cells (Figure 4- Figure supplemental 1B). Arguing that these functions likely depend on the presence of the CitA protein rather than citrate synthase enzymatic activity, the *citA* mutant phenotypes were not corrected by complementation of *citA* mutant cells with a multi-copy plasmid harboring *C. crescentus citB* (pMT335-*citB*) or *E. coli gltA* (pMT335-*gltA*) (Figure 4C). However, these deficiencies were corrected when a *WT* copy of *citA* was expressed *in trans* on a multi-copy plasmid (pMT335-*citA*) (Figure 4C). Thus, CitA promotes the G1àS transition, a function that other citrate synthases such as CitB and GltA cannot provide, despite having citrate synthase activity. Further support for the conclusion that CitA fulfills are regulatory role independent of catalytic activity came from discovering that catalytically inactive CitA still retained regulatory function. Residue H306 of *E. coli* GltA is critical to bind the oxaloacetate and its substitution prevents the catalytic activity of GltA (Pereira et al. 1994; Handford et al. 1988). We thus engineered variants in which the corresponding residue (H303) in *C. crescentus* CitA is substituted either by a tryptophan or an alanine, giving rise to the H303W and H303A CitA variants. As expected, expression of the CitA^H303W^ or CitA^H303A^ variant in *E. coli* Δ*gltA* cells no longer corrected the glutamate auxotrophy on minimal medium as determined by EOP assays (Figure 4- Figure supplemental 1C). Immunoblotting using polyclonal antibodies to CitA revealed that these variants are produced to the same levels as WT CitA (Figure 4- Figure supplemental 4D). We therefore conclude that CitA^H303W^ and CitA^H303A^ have lost enzymatic activity. When these variants were expressed in *C. crescentus* Δ*citA* mutant cells to similar levels than WT CitA (Figure 4- Figure supplemental 1E), a normal FACS profile and cell size distribution was observed by FACS and phase contrast microscopy, respectively (Figure 4D). In sum, these results show that the catalytic activity of CitA is dispensable for its cell cycle regulatory function.

**Figure 4.**
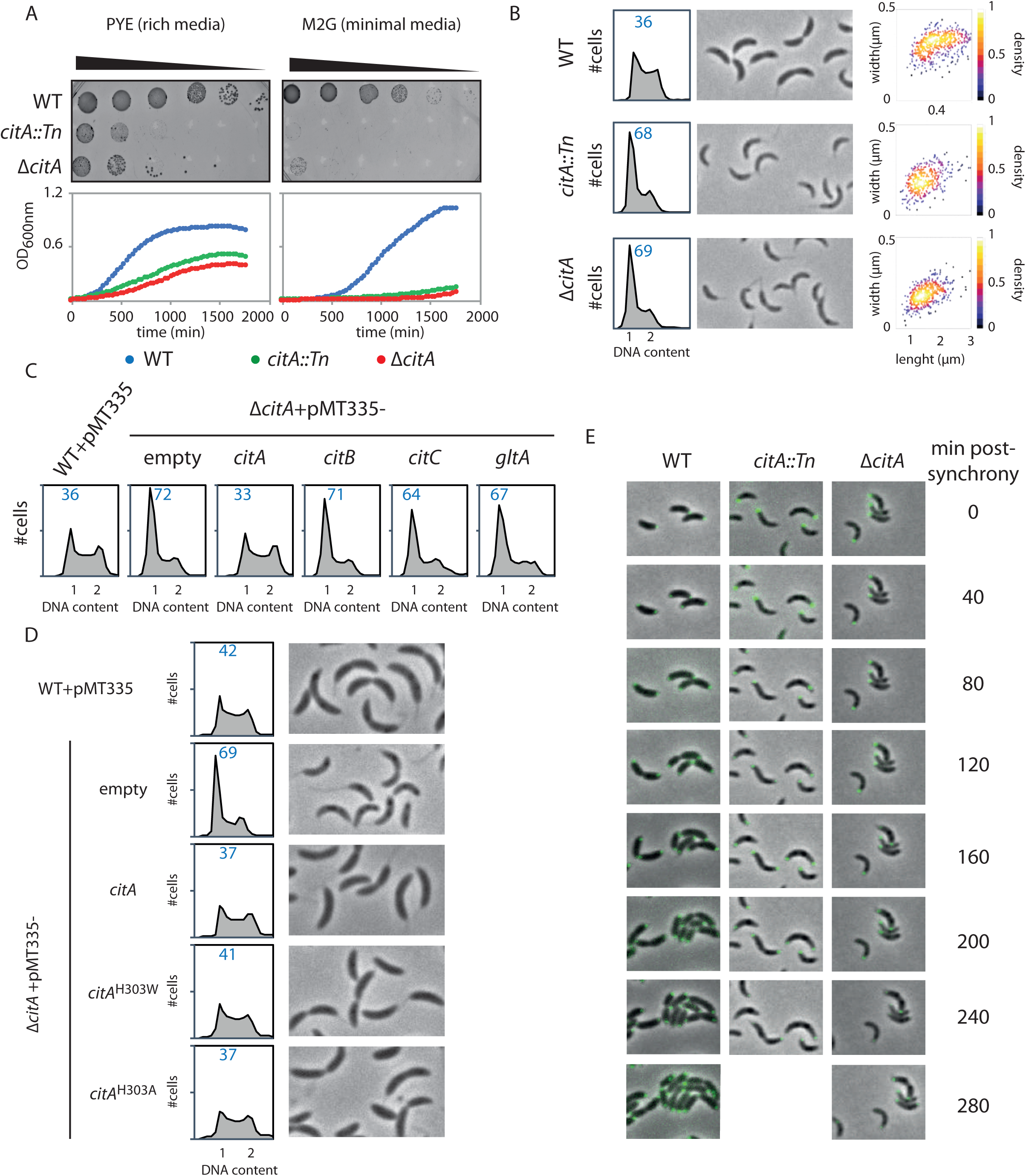
Inactivation of CitA induces a G1 block. (A) Spot dilution and growth curve of the WT (MB1), *citA::Tn* (MB2622) and Δ*citA* (MB2559). For spot dilution, cells were grown overnight in PYE, adjusted to OD_600nm_ ∼0.5 and serially diluted on a rich PYE medium (left upper part) or on a minimal M_2_G medium (right upper part). For growth curve, cells were grown overnight in PYE, washed twice with M_2_ buffer and similar amount of each strain was used to inoculate PYE medium (left bottom part) or M_2_G medium (right bottom part). (B) Flow cytometry profiles and phase contrast images of WT (MB1), *citA::Tn* (MB2622) and Δ*citA* (MB2559). Cells were exponentially grown in PYE and genome content was analyzed by FACS. Right part is a scatter plot of cell lengths and widths of each indicated population. (C) Flow cytometry profiles showing complementation of the Δ*citA* strain expressing an empty plasmid (MB3433) or *citA* (MB3435), *citB* (MB3469), *citC* (MB3471) from *C. crescentus* or the citrate synthase from *E. coli* (*gltA*) (MB3473). WT cells harboring an empty pMT335 are also shown (MB1537). (D) Flow cytometry profiles and phase contrast images of *C. crescentus* expressing a catalytic mutant of CitA. WT carrying an empty plasmid (MB1537), or Δ*citA* harboring an empty plasmid (MB3433) or citA (MB3435) or *citA*^H303A^ (MB3439) or *citA*^H303W^ (MB3437) are showed. (E) Time-lapse fluorescence microscopy of WT (MB557), *citA::Tn* (MB2452) and Δ*citA* (MB3467) harbouring a *parB*::*gfp*-*parB*. Cells were grown in PYE, synchronized and spotted on a PYE agarose pad. Each picture was taken every 20 minutes.

As the abundance of numerous regulators involved in the G1àS transition was previously shown to fluctuate in abundance during the cell cycle like SpmX (Radhakrishnan et al. 2008), KidO (Radhakrishnan et al. 2010), DivJ (Wheeler and Shapiro 1999) and CtrA (Domian et al. 1997), we wondered if the abundance of CitA is cell-cycle regulated. To this end, we monitored CitA abundance in synchronized cells by immunoblotting using polyclonal antibodies to CitA. In contrast to CtrA that is present in the swarmer and pre-divisional cells while absent in the stalked cell, CitA is present at a constant level along the cell cycle (Figure 4- Figure supplemental 1F), indicating that the cell cycle control function of CitA is mediated at the level of activity.

### CitA is required for S-phase entry in the presence of (p)ppGpp

To establish that CitA is required for the G1àS transition, cell cycle studies using synchronized *WT* and *citA* mutant cells were performed. FACS analysis revealed that *WT* cells initiate DNA replication 30 minutes after the release of G1 cells into PYE, whereas *citA*::Tn or *ΔcitA* cells do not enter S-phase before the 90 minute time point (Figure 4- Figure supplemental 1G). We also discovered that a fraction of *citA*::Tn or *ΔcitA* cells remained in G1 phase, with only approximately half entering S-phase. To confirm this interpretation, we conducted single cell time-lapse microscopy experiments with synchronized WT and *citA*::Tn or *ΔcitA* G1 cells expressing GFP-ParB as a marker for DNA replication (figure 4E). ParB is a chromosome partitioning protein that specifically binds near the origin of replication (C*_ori_*) and is translocated with a duplicated copy of C*_ori_* to the daughter cell compartment once DNA replication commences (Mohl and Gober 1997; Thanbichler and Shapiro 2008). In synchronized *WT* cells expressing ParB-GFP, we observed that G1 cells initially harbor a single, polarly localized C*_ori_*, represented by a single GFP-ParB focus. After 40 minutes, ∼80% (n=39) of the cells replicated C*_ori_* visualized as a duplicated GFP-ParB focus, one of which is segregated to the opposite pole. Finally, cell division is completed by 120 minutes. By contrast, in *citA*::Tn (n=35) or *ΔcitA* (n=29) cells, duplicated GFP-ParB foci only appeared in some cells 100 minutes after synchronization. Importantly, we noticed that even after 260 minutes, ∼60% of the population still exhibited only one GFP-ParB focus. Thus, a large fraction of the population remains in G1-phase and that only part of the *citA* mutant population enters S-phase.

To determine the genetic basis of the G1 block of *citA*::Tn or *ΔcitA* cells, we isolated fast growing suppressor mutants by serial dilution of two independent *citA*::Tn or Δ*citA* cultures, re-diluting them each day for 4 days (Figure 5A). Whole-genome sequencing of two *citA::*Tn and one Δ*citA* suppressor mutant that grew faster (identified as large colonies, Figure 5B) revealed a different frameshift mutation in the same region of the PEP-phosphotransferase domain-encoding region of PtsP (CCNA_00892) (Figure 5C). PtsP resembles the first enzyme of a nitrogen-related PEP-phosphotransferase (PTS^Ntr^) protein homologue (EI^Ntr^ in Enterobacteria) that typically uses PEP rather than ATP as the phospho-donor to phosphorylate clients proteins such as the HPr phospho-carrier protein (Deutscher et al. 2014) (Figure 5D). We hypothesized that the PtsP frameshift mutation in the *citA* suppressor mutants eliminated or decreased PtsP function. If so, an in-frame deletion in *ptsP* (Δ*ptsP*) should recapitulate the fast-growing phenotype of *citA* mutant cells. In agreement with this, when the Δ*citA* mutation was introduced into Δ*ptsP* cells, the resulting double mutants grew faster in PYE broth than the Δ*citA* single mutant (Figure 5- Figure supplemental 1A). Moreover, EOP assays of single and double mutants revealed that the Δ*ptsP* mutation increases the viability of *citA* mutant cells (Figure 5- Figure supplemental 1A). Finally, and importantly, the FACS profile of Δ*ptsP citA*::Tn double mutant cells mirrors that of *WT* cells, indicating that loss of PtsP indeed nullifies the severe G1 block caused by loss of CitA (Figure 5E).

**Figure 5.**
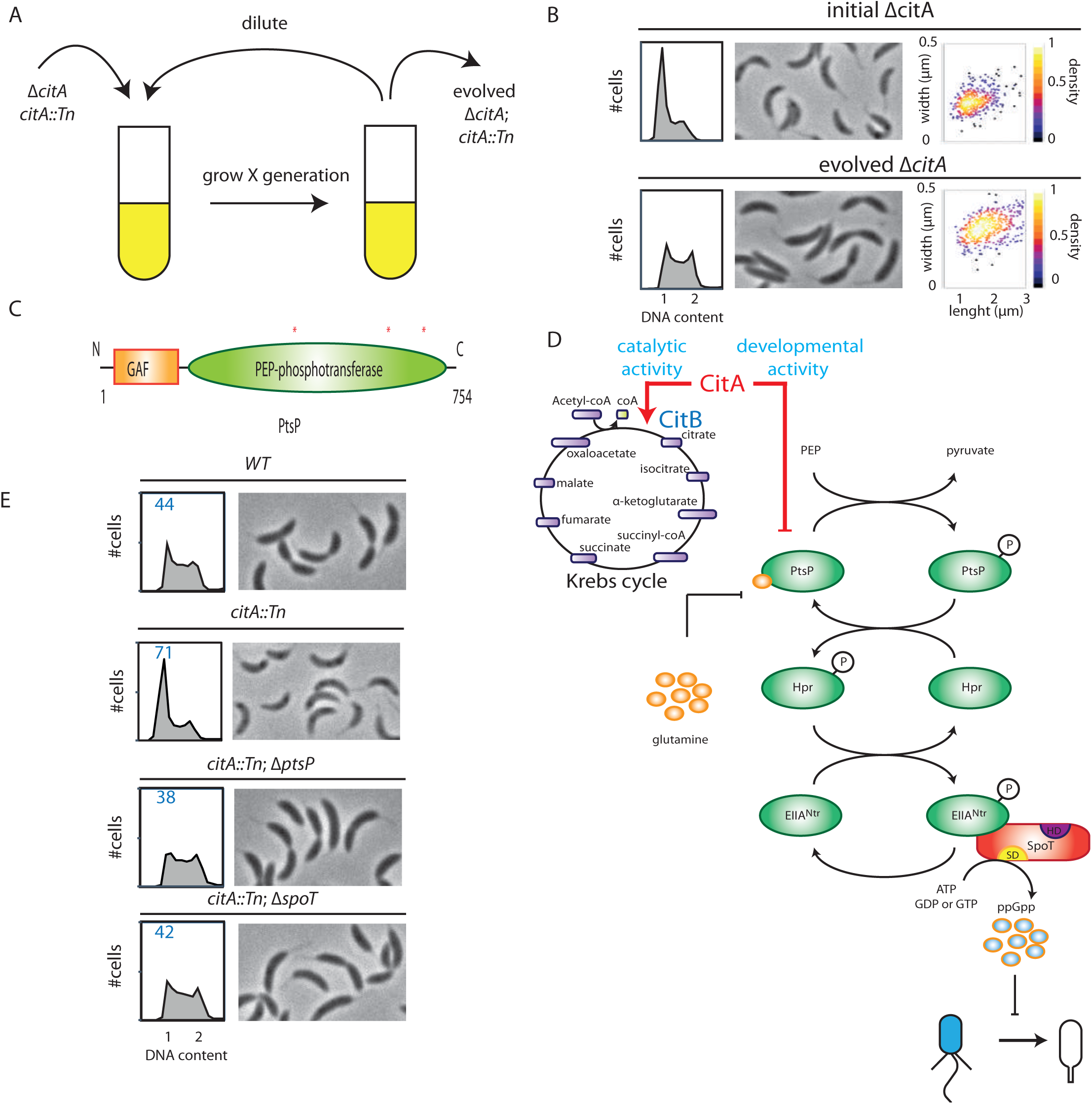
Absence of CitA induces (p)ppGpp production. (A) Cartoon of experimental evolution of Δ*citA* or *citA*::*Tn* by serial dilution. The suppressors were identified based on their ability to growth better in PYE medium after 4 days of dilution. (B) Flow cytometry profiles and phase contrast images of Δ*citA* (initial strain, upper part) or Δ*citA* after the evolution experiment (evolved strain, bottom part). Cells were exponentially grown in PYE and genome content was analyzed by FACS. Right part is a scatter plot of cell lengths and widths of each indicated population. (C) Domain organization of PtsP from the N to C terminus, indicating the total length in amino acid of the protein. The two domains, GAF and PEP-phosphotransferase are indicated. Asterisks indicate the position of the suppressive mutation leading to frameshift mutation in the PtsP PEP-phosphotransferase domain. (D)The Pts^Ntr^ pathway is represented (adapted from Ronneau et al. 2016). Intracellular glutamine regulates the autophosphorylation of PtsP. In case of nitrogen starvation, glutamine pool drops triggering PtsP phosphorylation leading to increase of phosphorylated EII^Ntr^. Once phosphorylated, EII^Ntr^ inhibits the hydrolase activity of SpoT leading to (p)ppGpp accumulation, blocking the swarmer to stalked cell transition. The two functions of CitA are represented, one acting as a metabolic enzyme into the Krebs cycle and the other one acting on the development of *C. crescentus*, independently of its catalytic activity. Absence of CitA activates the Pts^Ntr^ pathway, by a glutamine-independent mechanism, triggering (p)ppGpp production delaying the swarmer to stalk transition. (E) Flow cytometry profiles and phase contrast images of *WT* (MB1), *citA*::*Tn* (MB2622), Δ*spoT citA*::*Tn* (MB2413) and Δ*ptsP citA*::*Tn* (MB2426). Genome content was analyzed by FACS during exponential phase in PYE.

Since *C. crescentus* PtsP is known to inhibit the hydrolase activity of SpoT (Figure 5D), the bifunctional synthase/hydrolase of the (p)ppGpp alarmone that can extend the G1-phase of the cell cycle in *C. crescentus* (Stott et al. 2015; Gonzalez and Collier 2014; Ronneau et al. 2016), we reasoned that a Δ*spoT* mutation should also be epistatic to the *citA* mutation. Indeed, when the *citA*::Tn mutation was transduced into Δ*spoT* cells, the resulting double mutant exhibited a similar growth rate and FACS profile as the *WT* (Figure 5E, Figure 5- Figure supplemental 1A). In sum, these results suggest that the absence of *citA* leads to an activation of PtsP which in turn leads to an accumulation of (p)ppGpp in a SpoT-dependent manner. The resulting increase in (p)ppGpp then elicits the G1 arrest by maintaining CtrA activity. If true, then the *citA* mutation may have simply surfaced in the screen for Δ*tipN* Δ*cpdR* double mutant cells mutants with elevated CtrA activity because (p)ppGpp enhances CtrA activity. Consistent with this model, we found that the FACS profile is reversed in Δ*cpdR* Δ*tipN* Δ*citA* Δ*spoT* and Δ*cpdR* Δ*tipN* Δ*citA* Δ*ptsP* quadruple mutant compared to Δ*cpdR* Δ*tipN* Δ*citA* triple mutant cells, resembling that of the Δ*tipN* Δ*cpdR* double mutant cells (Figure 5- Figure supplemental 1B). These experiments show that the suppression by *citA* is dependent on the presence of PtsP and SpoT. Importantly, artificial induction of (p)ppGpp in Δ*tipN* Δ*cpdR* double mutant expression of a constitutively active form of the *E. coli* (p)ppGpp synthetase RelA (referred to RelA’) from the *C. crescentus xylX* locus (Gonzalez and Collier 2014) is sufficient to induce an increase in G1 cells (Figure 5- Figure supplemental 1C), whereas a catalytic inactive mutant of RelA’ (named RelA’^E335Q^) is unable to do so.

## DISCUSSION

To our surprise, a genetic selection for regulators of the G1-phase promoter, P*_pilA_*, in *C. crescentus* identified the gene encoding CitA citrate synthase as negative regulator of G1-phase. Our demonstration that CitA is a functional citrate synthase enzyme along with the fact that a catalytically active variant still retains the cell cycle control functions, implicates this protein as unprecedented coordinator of bacterial cell cycle progression and central (TCA) metabolism (Figure 5D).

Specifically, CitA promotes S-phase entry as evidenced by our finding that inactivation of CitA blocks the G1àS transition using (p)ppGpp (Boutte et al. 2012; Gonzalez and Collier 2014). Consistent with the notion that (p)ppGpp stalls cells in G1-phase by maintaining (active) CtrA, the *citA*::Tn mutation was isolated by a selection for elevated activity of the CtrA-dependent P*_pilA_* promoter. CtrA is regulated at the level of abundance through proteolysis (and synthesis) and at the level of activity by phosphorylation. Since CtrA is already rendered stable (by the *cpdR* mutation) in the mutant background in which the screen was conducted, our findings imply that the *citA*::Tn mutation and (p)ppGpp stimulates CtrA activity, a role of (p)ppGpp that had not be inferred in prior studies. With this effect on CtrA, arguably the master regulator of *C. crescentus* development, enzymatic activity of CitA is situated at key junction in central metabolism while fulfilling a key regulatory role in bacterial cell cycle control and differentiation, as suggested by the finding that nutritional stress may also act on CtrA in other alpha-proteobacteria such as the symbiont *Sinorhizobium meliloti* (De Nisco et al. 2014), raising the possibility that the ortholog (SMc02087, 70% identity to CitA) also contributes to cell cycle control and possibly plant symbiosis via (p)ppGpp or other effectors. As (p)ppGpp has also been implicated in regulating antibiotic tolerance in different species and recent links between TCA genes and antibiotic tolerance have been observed (Sinha et al.; Zalis et al. 2019), our finding that CitA mutants experience a (p)ppGpp-dependent G1 arrest, may provide an explanation of why bacteria are less susceptible to bactericidal antibiotics under TCA cycle stress.

### (p)ppGpp and metabolic control of the cell cycle

Remarkably, the cell cycle imbalances caused by the loss of CitA are mitigated when (p)ppGpp production is abolished (by the Δ*spoT* or Δ*ptsP* mutation), indicating that loss of the enzymatic activity of CitA is not detrimental to cells, whereas other TCA cycle enzymes are essential for viability. Interestingly, carbon starvation in *C. crescentus* leads to a phenotype resembling that of the G1-arrested Δ*citA* cells and a smaller cell volume induced by (p)ppGpp signaling (Lesley and Shapiro 2008; Leslie et al. 2015). Importantly, ectopic (p)ppGpp production from RelA’ phenocopies Δ*citA* cells, impairing the G1→S transition and retaining cells in the G1-phase swarmer (dispersal) state (Gonzalez and Collier 2014). The underlying mechanism is the production of the (p)ppGpp alarmone by the SpoT enzyme induced by PtsP. While PtsP has recently been shown to stimulate (p)ppGpp production in response to glutamine deprivation, thought to arise from nitrogen starvation (Ronneau et al. 2016), our metabolomic data reveal no difference in the glutamine pool comparing the WT and the Δ*citA*, indicating that CitA controls PtsP and SpoT via a different input. In support of this, addition of glutamine did not correct the defects of Δ*citA* cells. Indeed, the Pts^Ntr^ could be activated by several pathways leading to the (p)ppGpp production.

While (p)ppGpp signaling has mainly been described for cells experiencing nutritional stress conditions in stationary phase or in medium lacking nutrients, (p)ppGpp signaling has also been implicated in nutrient replete conditions, notably during the *C. crescentus* cell cycle (Boutte et al. 2012). In addition, a basal level of (p)ppGpp is crucial for global control of transcription, translation and cell size control in unstressed conditions in cyanobacteria (Puszynska and O’Shea 2017). A possible explanation for these effects is that cells experience nutritional stress during distinct cell cycle phases owing to metabolite fluctuations, caused by variabilities in enzyme abundance or activities during the cell cycle. Recently evidence has been provided that the cellular redox potential changes as a function of the *C. crescentus* cell cycle (Narayanan et al. 2015), suggesting the redox equivalents fluctuate. Since CitA is present throughout the cell cycle, it is possible that allosteric regulation by its substrates underlies the regulatory effects or other mechanisms or post-translational regulation may act on CitA.

Evidence supporting allosteric regulation has been provided for the cell cycle-regulated KidO protein, an NADH-binding oxidoreductase homolog, that is present in G1-phase and during cell constriction. KidO is a bifunctional enzyme that acts as a cell division inhibitor that binds FtsZ and it also acts negatively on the CtrA activation pathway (Radhakrishnan et al. 2010). These activities explain why abolishing the cell-regulated proteolysis of KidO by the CpdR-regulated ClpXP pathway can simultaneously shorten the G1-phase and impair cytokinesis. Inactivation of *kidO* in cells lacking TipN and CpdR alleviates these problems, showing that KidO is the major source of cellular mis-regulation in the absence of TipN, a polar organizer that marks the new cell pole and is re-localized to the division plane during constriction (Lam et al. 2006; Huitema et al. 2006). As TipN is also known to associate with late division proteins *in vivo* (Yeh et al. 2010; Goley et al. 2011), it is not surprising that cells lacking TipN are slightly filamentous and perhaps predisposed to accentuated cell division problems compared to *WT* cells when imbalances in cell division regulators such as KidO occur. Interestingly another division regulator that functions as moonlighting enzyme and that is degraded in a ClpXP and CpdR-dependent manner has been identified: the glutamate dehydrogenase GdhZ whose activity is modulated by glutamate and NADH (Beaufay et al. 2015). Together KidO and GdhZ show how substrate binding folds can be used to coordinate cell division in response to metabolic inputs, while CitA, independently of its enzymatic activity, links central metabolism with cell cycle development level through CtrA and (p)ppGpp.

### Enzyme redundancy and moonlighting

Expression of the paralog CitB from *C. crescentus* or the ortholog from *E. coli*, GltA, in Δ*citA* cells does not reverse the G1→S block either, even though both enzymes exhibit efficient citrate synthase activity in an *E. coli* reporter system that requires its activity for growth. The finding that addition of glutamine does not rescue the developmental problem of a Δ*citA* strain and that metabolite extractions from *citA* mutant cells do not reveal a major perturbance in the levels of TCAs, provide further support for the conclusion that the *citA* mutant phenotype is not simply caused by a metabolic deficiency of blocked citrate production. Rather, CitA is a moonlighting protein since that performs a regulatory function that is genetically separable from enzymatic activity.

The role of citrate synthase in development has been noted in other bacteria. *B. subtilis* cells lacking citrate synthase sporulate poorly (Ireton et al. 1995) and a citrate synthase mutant of *Streptomyces coelicolor* is unable to erect aerial mycelium (Viollier et al. 2001). Importantly, while the growth defect of the citrate synthase mutant in *S. coelicolor* on minimal medium was suppressed by glutamate, development remains perturbed. Thus, developmental events in bacteria may be controlled by switches and central metabolic enzymes serve as ideal checkpoint mechanisms that couple developmental gene expression to central metabolism. In this context, it is noteworthy that enolase in *E. coli* (Aït-Bara and Carpousis 2015) and aconitase in *C. crescentus* (Hardwick et al. 2011) are associated with the RNA degradosome, with the latter also fulfilling role as RNA binding protein, also in eukaryotic cells (Bandyra and Luisi 2018). It is also not surprising therefore that in certain organisms a multiplicity of such enzymes exists, possibly to permit specialization with different functions (e.g. in cell regulation) or to allow tailoring the metabolic (enzymatic) needs to specialized growth periods.

The viability of *C. crescentus ΔcitA* cells, whereas other TCA cycle enzymes seem to be indispensable for growth, supports our finding that the activity of a second citrate synthase isoform CitB can support TCA function in the absence of CitA. Since CitA and CitB are both functional as citrate synthases in a heterologous host such as *E. coli*, the genetic and metabolic framework for functional specialization of citrate synthases is provided in *C. crescentus*. In eukaryotic cells like *S. cerevisiae*, it was demonstrated the presence of several paralogs in different compartments of the cell. CIT1 is located in the mitochondria participating in the tricarboxylic acid cycle while CIT2 is located in the peroxisome acting in the glyoxylate cycle (Rosenkrantz et al. 1986; Kim et al. 1986). Interestingly, several bacterial genomes encode parologs of citrate synthase, notably in *B. subtilis* (Jin and Sonenshein 1994) or in *Pseudomonas aeruginosa* (Mitchell 1996). While the presence of several spatially regulated isoform make sense in eukaryotic cells, the absence of compartments in prokaryotes make unclear why bacteria encodes several citrate synthase, but could be linked with temporal functions such as a burst in TCA biosynthetic activity, however *a priori* it is not clear why this could not be achieved by dual promoter control. Therefore, the presence of paralogs is easiest to reconcile with functional specialization, that may have evolved from the same structural fold, perhaps exploiting substrate binding pockets. In this context, it is important to note that in *Podospora anserina*, a citrate synthase mutant strain exhibits a developmental phenotype impairing meiosis independently of its catalytic citrate synthase activity (Ruprich-Robert et al. 2002), reminiscent to our finding highlighting the citrate synthase as a key checkpoint in all kingdom life to co-ordinate cell development and metabolism.

## MATERIALS AND METHODS

### Strains and growth condition

Strains, plasmids and oligos are listed in Supplemental Table S3, S4 and S5. *C. crescentus* NA1000 (Marks et al. 2010) and derivatives were cultivated at 30°C in peptone yeast extract (PYE)-rich medium (2 g/L bactopeptone, 1 g/L yeast extract, 1 mM MgSO_4_, and 0.5 mM CaCl_2_) or in M2 minimal salts supplemented with 0.2% glucose (M2G, 0.87 g/L Na_2_HPO_4_, 0.54 g/L KH_2_PO_4_, 0.50 g/L NH_4_Cl, 0.2% [wt/vol] glucose, 0.5 mM MgSO_4_, 0.5 mM CaCl_2_, and 0.01 mM FeSO_4_) (Ely 1991). *E. coli* S17-1 *λpir* (Simon et al. 1983) and EC100D (Epicentre Technologies, Madison, WI) cells were grown at 37°C in Lysogeny Broth (LB)–rich medium (10 g/L NaCl, 5 g/L yeast extract, and 10 g/L tryptone). When appropriate, media were supplemented with antibiotics at the following concentrations (µg/mL in liquid/solid medium for *C. crescentus* strains; μg.mL^−1^in liquid/solid medium for *E. coli* strains): kanamycin (5/20 μg.mL^−1^; 20/20 μg.mL^−1^), tetracycline (1/1 μg mL^−1^; not appropriate), spectinomycin and streptomycin (in solid for *C. crescentus* only) (25/25, five respectively; 30/90 μg.mL^−1^), gentamycin (1/1; 10/25 μg.mL^−1^), aztreonam (in solid only) (2.5 μg.mL^−1^) and colistin (4 μg.mL^−1^). PYE plates containing 3% sucrose were used to select for loss of pNTPS138-derived plasmids by recombination when constructing mutants by double recombination. When needed, for *C. crescentus*, D-xylose was added at 0.3% final concentration, glucose at 0.2% final concentration. Glutamine was used at 9.3mM final in liquid and solid medium.

Swarmer cell isolation, electroporation, biparental mating (intergeneric conjugations) and bacteriophage fCr30-mediated generalized transduction were performed as described (Ely 1991) with slight modifications. Briefly, swarmer cells were isolated by Percoll density-gradient centrifugation at 4°C, followed by three washes and final re-suspension in pre-warmed (30°C) PYE. Electroporation was done from 1 mL overnight culture that had been washed three times in sterile water. Biparental mattings were done using exponential phase *E. coli* S17-1 donor cells and *C. crescentus* recipient cells washed in PYE and mixed at 1:3 ratio on a PYE plate. After 4–5 hours of incubation at 30°C, the mixture of cells was plated on PYE harboring aztreonam (to counter select *E. coli*) and the antibiotic that the conjugated plasmid confers resistance to. Generalized transductions using fCr30 were done by mixing 50 μL ultraviolet-inactivated fCr30 lysate with 500 μL stationary phase recipient cells, incubation for 2 hr, followed by plating on PYE containing antibiotic to select for the transduced DNA.

### Metabolite extraction

For metabolite extraction, *C. crescentus* were grown overnight at 30°C in PYE medium and diluted to reach an OD600nm∼0.4. 10 mL of cell culture were centrifuged at 2000g for 5 minutes at 4°C. Metabolism was then quenched by resuspending the pellet in 1 mL of precooled methanol/H_2_O (80:20 (vol/vol), kept at ∼ −20°C). Cells were subjected to lysis by five thaw/freeze (40°C/-80°C) cycles. Cellular debris was removed by centrifugation at 17,000g for 20 minutes at 4°C. Metabolite extracts were kept at −80°C prior to analysis on LC-MS. Bacterial biomass of individual samples was determined for normalization. The supernatants were completely evaporated using a SpeedVac (ThermoFisher, Langenselbold, Germany) and metabolite extracts were reconstituted in 100 µL acetonitrile:H_2_O 50:50. Quality control (QC) and diluted QC (dQC, diluted by 50%) samples were prepared by pooling equivalent volumes of all reconstituted samples and injected at a regular interval of 5 samples to assess analytical variability.

### Liquid Chromatography-High Resolution Mass Spectrometry (LC-HRMS) analysis

LC experiments were performed on a Waters H-Class Acquity UPLC system composed of a quaternary pump, an auto-sampler including a 15 μL flow-through-needle injector and a two-way column manager (Waters, Milford, USA) for which temperatures were set at 7 °C and 40°C respectively. The injected volume was 10 μL. Samples were analyzed with a hydrophilic liquid interaction chromatography (HILIC) SeQuant Zic-pHILIC column (150 x 2.1 mm, 5 μm) and the appropriate guard kit. For mobile phases, solvent A was acetonitrile and solvent B was H_2_O containing 2.8 mM ammonium formate adjusted at pH 9.00. Column flow rate was set at 300 µL.min^-1^. The following gradient was applied: 5% B for one minute, increased to 51% B over 9 minutes, holding for 3 minutes at 51% B and then returning back to 5% B in 0.1 minutes and re-equilibrating the column for 6.9 minutes. The UPLC system was coupled to a TWIMS-QTOF high resolution HRMS (Vion, Waters, Manchester, UK) through an electrospray ionization (ESI) interface. Analyses were performed in negative ESI mode and continuum data in the range of 50 - 1000 m/z were acquired with a scan time of 0.2 seconds. The ESI parameters were respectively set as follows: capillary voltage was −2.0 kV, source and desolvation temperatures were set at 120 and 500 °C, cone and desolvation gas flow were 50 and 800 L/h. Velocity and height of StepWave1 and StepWave2 were set to 300 m/s and 5 V and to 200 m/s and 30 V, respectively. The high definition MS^E^ (HDMS^E^, using ion mobility) settings consisted of trap wave velocity at 100 m/s; trap pulse height A at 10 V; trap pulse height B at 5 V; IMS wave velocity at 250 m/s; IMS pulse height at 45 V; wave delay set at 20 pushes and gate delay at 0 m/s. Gas flows of ion mobility instrument were set to 1.60 L/minute for trap gas, and 25 mL/min for IMS gas. Buffer gas was nitrogen. Fragmentation was performed in HDMS^E^ mode. For the collision energy, 6.0 eV was used for low energy and high energy was a ramp from 10 to 60 eV. Nitrogen was used as collision gas. Leucine-encephalin served as a lock-mass (554.2615 m/z for ESI-) infused at 5-minute intervals. The CCS and mass calibration of the instrument were done with the calibration mix “Major mix IMS-TOF calibration” (Waters, Manchester, UK). UNIFI v1.9.3 was used for data acquisition and data treatment.

### Analysis of raw LC-MS data

Run alignment, peak picking, adduct deconvolution and feature annotation were sequentially performed on Progenesis QI v2.3 (Nonlinear Dynamics, Waters, Newcastle upon Tyne, UK). Detected peaks were annotated with regard to a set of pure reference standards (MSMLS Library of Standards, Sigma-Aldrich) measured under the same experimental conditions as described elsewhere (Pezzatti et al. 2019b). The following tolerances were used: 2.5 ppm for precursor and fragment mass, 10% for retention time (Rt) and 5 % in the case of collisional cross section (CCS). Data processing was achieved by SUPreMe, an in-house software with capabilities for drift correction, noise filtering and sample normalization. Finally, data were transferred to SIMCA-P 15.0 software (Umetrics, Umea, Sweden) for multi-variate analysis (MVA).

### Microscopy and image analysis

Exponential phase *C. crescentus* cells cultivated in PYE were immobilized on a thin layer of 1.2% agarose. For *C. crescentus* time-lapse experiments, cells were first synchronized by Percoll density-gradient centrifugation and then immobilized on a thin layer of 1.2% agarose in PYE. Fluorescence and contrast microscopy images were taken with a phase contrast objective (Zeiss, alpha plan achromatic 100X/1.46 oil phase 3) on an Axio Imager M2 microscope (Zeiss) with appropriate filter (Visitron Systems GmbH) and a cooled CCD camera (Photometrics, CoolSNAP HQ2) controlled through Metamorph (Molecular Devices). Images were acquired and processed with ImageJ via Fiji software (Schneider et al. 2012; Schindelin et al. 2012). To perform cell segmentation and tracking, images were processed using MicrobeJ (Ducret et al. 2016). Statistics were performed on experiments performed in triplicate representing more than 300 cells.

### Genome-wide transposon mutagenesis coupled to deep-sequencing (Tn-Seq)

Pools of >100,000 Tn mutants were isolated as kanamycin-aztreonam or kanamycin-colistin resistant clones in the NA1000 (WT), Δ*tipN, ΔcpdR*::Ω backgrounds, with the same protocol as previously described using a mini-*himar1* Tn encoding kanamycin resistance (Viollier et al. 2004). For each Tn pool, chromosomal DNA was extracted and used to generate a Tn-Seq library sequenced on an Illumina HiSeq 2500 sequencer (Fasteris, Geneva, Switzerland). The single-end sequence reads (50 bp) stored in FastQ files were mapped against the genome of *Caulobacter crescentus* NA1000 (NC_011916) (Marks et al. 2010) genome and converted to BED files using BWA-MEM and bedtools BAM to BED tools respectively from the Galaxy server (https://usegalaxy.org/). The resulting BED file was imported into SeqMonk (http://www.bioinformatics.babraham.ac.uk/projects/seqmonk/) to build sequence read profiles. The initial quantification of the sequencing data was done in SeqMonk: the genome was subdivided into 50 bp probes, and for every probe we calculated a value that represents a normalized read number per million. A ratio for each 50bp position was done between the reads obtained in the Δ*tipN* or Δ*cpdR* strains to the *WT* reads. This file was used to generate the zoomed panels of the *popA*, *rcdA* and *cpdR* loci (Figure 1B) or the *tipN* locus (Figure 1- Figure supplemental 1A and 1B).

### Identification of *citA* (pilA-nptII suppressors)

The *citA*::Tn insertion was identified using a modification of the kanamycin resistance suppressor screen (Radhakrishnan et al. 2010). Briefly, we screened for mini-*himar1* Tn insertions that restore P*_pilA_* firing to Δ*tipN* Δ*cpdR* double mutant cells harboring the P*_pilA_*-*nptII* transcriptional reporter that confers kanamycin resistance to 20 μg ml^−1^ when P*pilA* is fully active. The Tn encodes gentamycin resistance on plasmid pMar2xT7 was delivered from *E. coli* S17-1 *λpir* (Liberati et al. 2006) to Δ*tipN* Δ*cpdR pilA*::P*_pilA_*-*nptII C. crescentus* cells by selected on plates gentamycin (1 μg ml^−1^), kanamycin (20 μg ml^−1^) and aztreonam (2.5 μg ml^−1^, to counter-select *E. coli*). This screen gave rise to one isolate Φ40 with the desired resistance profile. The Tn insertion in Φ40was mapped to the uncharacterized *CCNA_01983* gene at nucleotide (nt) position 1061847 of the *C. crescentus* NA1000 genome sequence using arbitrarily primed PCR (Liberati et al. 2006).

### Evolution experiment

Two independent clones freshly transduced *C. crescentus* NA1000 with Δ*citA*::*kan* or *citA*::Tn were inoculated in 3mL of PYE. Stationary phase cultures were diluted in 3 mL PYE to an optical density OD_600nm_∼0.02. After 2 days, the 4 cultures were re-diluted to an OD_600nm_ ∼0.001 in 3 mL PYE. The phenotype of each strain was checked by phase contrast microscopy and FACS analysis. Each culture was streaked on a PYE plate and one single colony from each culture was grown overnight and chromosomal DNA was extracted. Three suppressors were subjected to whole-genome sequencing. Library preparation and sequencing were performed by the Genomic platform iGE3 at the university of Geneva on a HiSeq 2500 with 50bp paired-end reads. Data analysis to identify mutations was done using freebayes v1.1.0-3 (Garrison and Marth 2012) against the *C. crescentus* NA1000 reference genome (NC_011916.1).

### Growth curve

The overnight cultures were started in PYE or in M2G. The cultures were diluted to obtain an OD_600nm_ of 0.1 in PYE or M2G and were incubated at 30°C with a continuous shaking in a microplate reader (Synergy H1, Biotek). The OD_600nm_ was recorded every 30 minutes for 30 hours. The graph represents the trend of the growth curve of three independent experiments.

### Fluorescence-activated cell sorting (FACS)

Cells in exponential growth phase (OD_600_, 0.3 to 0.6) were fixed 1:10 (vol/vol) in ice-cold 70% ethanol solution and stored at −20 °C until further use. For rifampicin treatment, the mid-log phase cells were grown in the presence of 20 µg/mL rifampicin at 30°C for 3 hours. Cells were fixed as mentioned above. Fixed cells were centrifuged at 6200g for 3 minutes at room temperature and washed once in FACS staining buffer (pH 7.2; 10 mM Tris-HCl, 1 mM EDTA, 50 mM Na-citrate, 0.01% Triton X-100). Then, cells were centrifuged at 6200g for 3 minutes at room temperature, resuspended in FACS staining buffer containing RNase A (Roche) at 0.1 mg.mL^-1^ for 30 minutes at room temperature. Cells were stained in FACS staining buffer containing 0.5 µM of SYTOX green nucleic acid stain solution (Invitrogen) and then analyzed using a BD Accuri C6 flow cytometer instrument (BD Biosciences, San Jose, CA, United States). Flow cytometry data were acquired and analyzed using the CFlow Plus v1.0.264.15 software (Accuri Cytometers Inc.). A total of 20,000 cells were analyzed from each biological sample, performed in triplicates. The green fluorescence (FL1-A) parameters was used to determine cell chromosome contents. Flow cytometry profiles within one figure were recorded in the same experiment, on the same day with the same settings. The scales of y- and x-axes of the histograms within one figure panel are identical. Each experiment was repeated independently three times and representative results are shown. The relative chromosome number was directly estimated from the FL1-A value of NA1000 cells treated with 20 µg/mL rifampicin for 3 hours at 30°C. Rifampicin treatment of cells blocks the initiation of chromosomal replication but allows ongoing rounds of replication to finish.

### Whole-cell extracts preparation

Five hundred μL of an exponential *Caulobacter* or *E. coli* cells (OD_600nm_ = 0.4 and 0.8 respectively) were harvested with 20,000g at 4°C for 5 minutes. Whole-cell extracts were prepared by resuspension of cell pellets in 75 µL TE buffer (10 mM Tris-HCl pH 8.0 and 1 mM EDTA) followed by addition of 75 µL loading buffer 2X (0.25 M Tris pH 6.8, 6% (wt/vol) SDS, 10 mM EDTA, 20% (vol/vol) Glycerol) containing 10% (vol/vol) β-mercaptoethanol. Samples were normalized for equivalent loading using OD_600nm_ and were heated for 10 minutes at 90°C prior to loading.

### Immunoblot analysis

Protein samples were separated by SDS–polyacrylamide gel electrophoresis and blotted on polyvinylidenfluoride membranes (Merck Millipore). Membranes were blocked overnight with Tris-buffered saline 1X (TBS) (50 mM Tris-HCl, 150 mM NaCl, pH 8) containing, 0.1% Tween-20 and 8% dry milk and then incubated for an additional three hours with the primary antibodies diluted in TBS 1X, 0.1% Tween-20, 5% dry milk. The different polyclonal antisera to CitA (1:5,000), CtrA (1:5,000) were used. Primary antibodies were detected using HRP-conjugated donkey anti-rabbit antibody (Jackson ImmunoResearch) with ECL Western Blotting Detection System (GE Healthcare) and a luminescent image analyzer (Chemidoc^Tm^ MP, Biorad).

### CitA purification and production of antibodies

Recombinant CitA protein was expressed as an N-terminally His_6_-tagged variant from pET28a in *E. coli* BL21(DE3)/ pLysS and purified under native conditions using Ni^2+^ chelate chromatography. Cells were grown in LB at 37°C to an OD_600nm_ of 0.6 and induced by the addition of IPTG to 1 mM for 3 hours and harvested at 5000 RPM at 4°C for 30 minutes. Cells were pelleted and re-suspended in 25 mL of lysis buffer (10 mM Tris HCl (pH 8), 0.1 M NaCl, 1.0 mM β-mercaptoethanol, 5% glycerol, 0.5 mM imidazole Triton X-100 0.02%). Cells were sonicated in a water–ice bath, 15 cycles of 30 seconds ON; 30 seconds OFF. After centrifugation at 5000g for 20 minutes at 4°C, the supernatant was loaded onto a column containing 5 mL of Ni-NTA agarose resin (Qiagen, Hilden, Germany) pre-equilibrated with lysis buffer. The column was rinsed with lysis buffer, 400 mM NaCl and 10 mM imidazole, both prepared in lysis buffer. Fractions were collected (in 300 mM Imidazole buffer, prepared in lysis buffer) and used to immunize New Zealand white rabbits (Josman LLC).

### Strain construction

#### MB3075 (NA1000 Δ*tipN* Δ*popA*)

A pNTPS138 derivative (pNTPS138-Δ*tipN*) (Huitema et al. 2006) was integrated nearby the marker-less Δ*tipN* mutation by homologous recombination. Phage fCr-30-mediated generalized transduction was used to transfer the mutant Δ*tipN* allele into the recipients NA1000 Δ*popA* by selecting for kanamycin resistance. Clones that have lost pNPTS138-*ΔtipN* by homologous recombination were probed for kanamycin resistance (on PYE plates supplemented with kanamycin) following sucrose counter-selection. PCR was used to verify the integrity of the mutants.

#### MB3079 (NA1000 Δ*tipN* Δ*rcdA::*Ω)

A pNTPS138 derivative (pNTPS138-Δ*tipN*) (Huitema et al. 2006) was integrated nearby the marker-less Δ*tipN* mutation by homologous recombination. Phage fCr-30-mediated generalized transduction was used to transfer the mutant Δ*tipN* allele into the recipients NA1000; Δ*rcdA::*Ω by selecting for kanamycin resistance. Clones that have lost pNPTS138-*ΔtipN* by homologous recombination were probed for kanamycin resistance (on PYE plates supplemented with kanamycin) following sucrose counter-selection. PCR was used to verify the integrity of the mutants.

#### MB2017 (NA1000 Δ*tipN* Δ*cpdR::tet*)

The Δ*cpdR::tet* allele was introduced into NA1000 Δ*tipN* by generalized transduction using ϕCr30 and then selected on PYE plates containing tetracycline.

#### MB2366 (NA1000 Δ*tipN* xylX::kidO^AA::DD^)

The *xylX*::*kidO*^AA::DD^ (kan^R^) allele was introduced into NA1000 Δ*tipN* by generalized transduction using ϕCr30 and then selected on PYE plates containing kanamycin.

#### MB2720 (NA1000 Δ*tipN* Δ*cpdR::tet* Δ*kidO*)

A pNTPS138 derivative (pNTPS138-Δ*tipN*) (Huitema et al. 2006) was integrated nearby the marker-less Δ*tipN* mutation by homologous recombination. ϕCr-30-mediated generalized transduction was used to transfer the mutant Δ*tipN* allele into the recipients NA1000 Δ*kidO* by selecting for kanamycin resistance. Clones that have lost pNPTS138-*ΔtipN* by homologous recombination were probed for kanamycin resistance (on PYE plates supplemented with kanamycin) following sucrose counter-selection. PCR was used to verify the integrity of the mutants. Then, Δ*cpdR::tet* allele was introduced into NA1000 Δ*tipN* Δ*kidO* by transduction using ϕCr30 and then selected on PYE plates containing tetracycline.

#### MB2325 (NA1000 *pilA*::P*_pilA_*-*GFP*)

The *pilA*::P*_pilA_*-*GFP* (kan^R^) allele was introduced into NA1000 by generalized transduction using ϕCr30 and then selected on PYE plates containing kanamycin.

#### MB2327 (NA1000 Δ*cpdR::*Ω *pilA*::P*_pilA_*-*GFP*)

The *pilA*::P*_pilA_*-*GFP* (kan^R^) allele was introduced into NA1000 Δ*cpdR::*Ω (Spc^R^) by generalized transduction using ϕCr30 and then selected on PYE plates containing kanamycin.

#### MB2329 (NA1000 Δ*tipN pilA*::P*_pilA_*-*GFP*)

The *pilA*::P*_pilA_*-*GFP* (kan^R^) allele was introduced into NA1000 Δ*tipN* by generalized transduction using ϕCr30 and then selected on PYE plates containing kanamycin.

#### MB2331 (NA1000 Δ*tipN* Δ*cpdR::*Ω *pilA*::P*_pilA_*-*GFP*)

The *pilA*::P*_pilA_*-*GFP* (kan^R^) allele was introduced into MB2017 (NA1000 Δ*tipN* Δ*cpdR::*Ω by generalized transduction using ϕCr30 and then plated on PYE containing kanamycin.

#### MB2268 (NA1000 *pilA*::P*_pilA_*-*nptII*)

The *pilA*::P*_pilA_*-*nptII* (Spc^R^) allele was introduced into NA1000 by generalized transduction using ϕCr30 and then selected on PYE plates containing spectinomycin.

#### MB2271 (NA1000 ΔtipN ΔcpdR::tet pilA::P_pilA_-nptII)

The *pilA*::P*_pilA_*-*nptII* (Spc^R^) allele was introduced into MB2017 (NA1000 Δ*tipN* Δ*cpdR::tet*) by generalized transduction using ϕCr30 and then selected on PYE plates containing spectinomycin.

#### MB2559 (NA1000 *ΔcitA*::pNTPS138ΔcitA)

A pNTPS138 derivative (pNTPS138-Δ*citA*) was integrated nearby the marker-less Δ*citA* mutation by homologous recombination. ϕCr-30-mediated generalized transduction was used to transfer the mutant Δ*citA* allele into the recipients NA1000 by selecting for kanamycin resistance on PYE plates containing kanamycin.

#### MB3056 (NA1000 Δt*ipN ΔcpdR::tet citA::Tn pilA::*P*_pilA_-nptII*)

The *citA*::Tn (Gent^R^) allele was introduced into MB2271 (NA1000 Δ*tipN* Δ*cpdR::tet pilA*::P*_pilA_*-*nptII*) cells by transduction using ϕCr30 and then selected on PYE plates containing gentamycin.

#### MB3058 (NA1000 Δ*tipN ΔcpdR::tet ΔcitA pilA::*P*_pilA_-nptII*)

ϕCr-30-mediated generalized transduction was used to transfer the mutant Δ*citA* allele from MB2559 into MB2017 (NA1000; Δ*tipN* Δ*cpdR::tet*) recipient cells by selecting for kanamycin resistance. Clones that have lost pNPTS138-*ΔcitA* by homologous recombination were probed for kanamycin resistance (on PYE plates supplemented with kanamycin) following sucrose counter-selection (giving rise to strain named MB3054. PCR was used to verify the integrity of the mutants. Then, the *pilA*::P*_pilA_*-*nptII* (Spc^R^) allele was introduced into MB3054 (NA1000 Δ*tipN* Δ*cpdR::tet* Δ*citA*) by generalized transduction using ϕCr30, selecting on PYE plates containing spectinomycin.

#### MB2679 (NA1000 Δ*citBC*)

The markerless Δ*citBC* double mutant was created by introducing into the *WT* (NA1000) using the standard two-step recombination sucrose counter-selection procedure induced by the pNTPS138-Δ*citBC* (pMB309). Briefly, first integration was done by matting of the eMB552 (S17-1 carrying the pMB309) and *C. crescentus* NA1000, selecting for kanamycin and aztreonam (to eliminate the donor strain). Clones that have lost pNPTS138-*ΔtipN* by homologous recombination were probed for kanamycin resistance (on PYE plates supplemented with kanamycin) following sucrose counter-selection (giving rise to strain named MB2679. PCR, using outside primers that do not hybridize within the Δ*citBC* deletion carried on pNTPS138, was used to verify the integrity of the mutants.

#### MB2622 (NA1000 *citA*::Tn)

The *citA*::Tn (Gent^R^) allele was introduced into NA1000 by generalized transduction using ϕCr30 and then selected on PYE plates containing gentamycin.

#### MB1537 (NA1000; pMT335)

Plasmid pMT335 was introduced into NA1000 by electroporation and then plated on PYE harboring gentamycin.

#### MB3433 (NA1000 Δ*citA*; pMT335)

ϕCr-30-mediated generalized transduction was used to transfer the mutant Δ*citA* allele from MB2559 into MB1537 recipient cells by selecting for kanamycin resistance.

#### MB3435 (NA1000 Δ*citA*; pMT335*-citA*)

Plasmid pMB302 (pMT335-*citA*) was introduced into NA1000 by electroporation and then plated on PYE harboring gentamycin. ϕCr-30-mediated generalized transduction was used to transfer the mutant Δ*citA* allele from MB2559 into NA1000; pMT335-*citA* cells by selecting for kanamycin resistance.

#### MB3469 (NA1000 Δ*citA*; pMT335*-citB*)

Plasmid pMB303 (pMT335-*citB*) was introduced into NA1000 by electroporation and then plated on PYE harboring gentamycin. ϕCr-30-mediated generalized transduction was used to transfer the mutant Δ*citA* allele from MB2559 into NA1000; pMT335-*citB* cells by selecting for kanamycin resistance.

#### MB3471 (NA1000 Δ*citA*; pMT335*-citC*)

Plasmid pMB304 (pMT335-*citC*) was introduced into NA1000 by electroporation and then plated on PYE harboring gentamycin. ϕCr-30-mediated generalized transduction was used to transfer the mutant Δ*citA* allele from MB2559 into NA1000; pMT335-*citC* cells by selecting for kanamycin resistance.

#### MB3473 (NA1000 Δ*citA*; pMT335*-gltA*)

Plasmid pMB310 (pMT335-*gltA*) was introduced into NA1000 by electroporation and then plated on PYE harboring gentamycin. ϕCr-30-mediated generalized transduction was used to transfer the mutant Δ*citA* allele from MB2559 into NA1000; pMT335-*gltA* cells by selecting for kanamycin resistance.

#### MB3437 (NA1000 Δ*citA*; pMT335*-citA^H303W^*)

Plasmid pMB325 (pMT335-*citA^H303W^*) was introduced into NA1000 by electroporation and then plated on PYE harboring gentamycin. ϕCr-30-mediated generalized transduction was used to transfer the mutant Δ*citA* allele from MB2559 into NA1000; pMT335*-citA^H303W^* cells by selecting for kanamycin resistance.

#### MB3439 (NA1000 Δ*citA*; pMT335*-citA^H303A^*)

Plasmid pMB326 (pMT335-*citA^H303A^*) was introduced into NA1000 by electroporation and then plated on PYE harboring gentamycin. ϕCr-30-mediated generalized transduction was used to transfer the mutant Δ*citA* allele from MB2559 into NA1000; pMT335*-citA^H303A^* cells by selecting for kanamycin resistance.

#### MB2452 (NA1000 *parB*::*GFP*-*parB citA*::Tn)

The *citA*::Tn (Gent^R^) allele was introduced into MB557 (NA1000; *parB*::*GFP*-*parB*) by generalized transduction using ϕCr30 and then plated on PYE plates containing gentamycin.

#### MB3467 (NA1000 *parB*::*GFP*-*parB* Δ*citA*)

ϕCr-30-mediated generalized transduction was used to transfer the mutant Δ*citA* allele from MB2559 into MB557 (NA1000; *parB*::*GFP*-*parB*) by selecting for kanamycin resistance on plates containing kanamycin.

#### MB2413 (NA1000 Δ*spoT citA*::Tn)

ϕCr-30-mediated generalized transduction was used to transfer the *citA*::Tn allele into MB2403 (NA1000 Δ*spoT*) cells by selection on plates PYE containing gentamycin.

#### MB2426 (NA1000 Δ*ptsP citA*::Tn)

ϕCr-30-mediated generalized transduction was used to transfer the *citA*::Tn allele into MB2417 (NA1000 Δ*ptsP*) cells by selectiion on plates PYE containing gentamycin.

#### MB3382 (NA1000 Δ*tipN ΔcpdR::tet ΔspoT ΔcitA)*

A pNTPS138 derivative (pNTPS138-Δ*spoT*) was integrated nearby the marker-less Δ*spoT* mutation by homologous recombination. Then, ϕCr-30-mediated generalized transduction was used to transfer the mutant Δ*spoT* allele into NA1000 Δ*tipN* cells by selecting for kanamycin resistance. Clones that have lost pNPTS138-*ΔspoT* by homologous recombination were probed for kanamycin resistance (on PYE plates supplemented with kanamycin) following sucrose counter-selection. PCR was used to verify the integrity of the mutants. ϕCr30-mediated generalized transduction was then used to transfer the mutant Δ*citA* allele from MB2559 into NA1000 Δ*tipN* Δ*spoT* recipient cells by selecting for kanamycin resistance. Finally, Δ*cpdR::tet* allele was introduced into NA1000 Δ*tipN* Δ*spoT* Δ*citA* recipient cells by transduction using ϕCr30, followed by selection on PYE plates containing tetracycline.

#### MB3386 (NA1000 Δ*tipN ΔcpdR::tet ΔspoT ΔcitA*)

A pNTPS138 derivative (pNTPS138-Δ*ptsP*) was integrated nearby the marker-less Δ*ptsP* mutation by homologous recombination. ϕCr-30-mediated generalized transduction was used to transfer the mutant Δ*ptsP* allele into the recipients NA1000 Δ*tipN* by selecting for kanamycin resistance. Clones that have lost pNPTS138-*ΔptsP* by homologous recombination were probed for kanamycin resistance (on PYE plates supplemented with kanamycin) following sucrose counter-selection. PCR was used to verify the integrity of the mutants. A pNTPS138 derivative (pNTPS138-Δ*citA*) was integrated nearby the marker-less Δ*citA* mutation by homologous recombination. ϕCr-30-mediated generalized transduction was then used to transfer the mutant Δ*citA* allele into NA1000 Δ*tipN* Δ*ptsP* recipient cells by selecting for kanamycin resistance. No counterselection was done. Finally, Δ*cpdR::tet* allele was introduced into NA1000 Δ*tipN* Δ*ptsP* Δ*citA* by transduction using ϕCr30, followed by selection on PYE plates containing tetracycline.

#### MB3366 (NA1000 Δ*tipN ΔcpdR::tet xylX::relA’-flag*)

The *xylX*::*relA’* (GentR) allele was introduced into NA1000 Δ*tipN*; by transduction using ϕCr30 and then plated on PYE harboring gentamycin. Then, Δ*cpdR::tet* allele was introduced into NA1000; Δ*tipN xylX*::*relA’* by transduction using ϕCr30 and then plated on PYE containing tetracycline.

#### MB3368 (NA1000 Δ*tipN ΔcpdR::tet xylX::relA’-flag*)

The *xylX*::*relA’*^E335Q^ (GentR) allele was introduced into NA1000 Δ*tipN* cells by transduction using ϕCr30, selected PYE plates containing gentamycin. Then, the Δ*cpdR::tet* allele was introduced into NA1000 Δ*tipN xylX*::*relA’*^E335Q^ recipient cells by transduction using ϕCr30, selecting on PYE plates containing tetracycline.

#### eMB554 (BW35113; pMT335)

Plasmid pMT335 was introduced into BW35113 by electroporation and then plated on LB agar containing gentamycin.

#### eMB556 (BW35113; Δ*gltA*::770; pMT335)

Plasmid pMT335 was introduced into JW0710-1 (BW35113; *ΔgltA770*::*kan*) by electroporation and then plated on LB agar containing gentamycin.

#### eMB558 (BW35113; Δ*gltA*::770; pMT335-*citA*)

Plasmid pMB302 (pMT335-*citA*) was introduced into JW0710-1 (BW35113; *ΔgltA770*::*kan*) by electroporation and then plated on LB agar containing gentamycin.

#### eMB560 (BW35113; Δ*gltA*::770; pMT335-*citB*)

Plasmid pMB303 (pMT335-*citB*) was introduced into JW0710-1 (BW35113; Δ*gltA770*::*kan*) by electroporation and then plated on LB agar containing gentamycin.

#### eMB562 (BW35113; Δ*gltA*::770; pMT335-*citC*)

Plasmid pMB304 (pMT335-*citC*) was introduced into JW0710-1 (BW35113; Δ*gltA770*::*kan*) by electroporation and then plated on LB agar containing gentamycin.

#### eMB564 (BW35113; Δ*gltA*::770; pMT335-*gltA*)

Plasmid pMB310 (pMT335-*gltA*) was introduced into JW0710-1 (BW35113; Δ*gltA770*::*kan*) by electroporation and then plated on LB agar containing gentamycin.

#### eMB581 (BW35113; Δ*gltA*::770; pMT335-*citA^H303W^)*

Plasmid pMB325 (pMT335-*citA^H303W^*) was introduced into JW0710-1 (BW35113; Δ*gltA770*::*kan*) by electroporation and then plated on LB agar containing gentamycin.

#### eMB581 (BW35113; Δ*gltA*::770; pMT335-*citA^D361E^)*

Plasmid pMB327 (pMT335-*citA^D361E^*) was introduced into JW0710-1 (BW35113; Δ*gltA770*::*kan*) by electroporation and then plated on LB agar containing gentamycin.

### Plasmid constructions

#### pMB278 (pNTPS138-Δ*citA*)

The plasmid construct used to delete *citA* (*CCNA_01983*) was made by PCR amplification of two fragments. The first to amplify the upstream region of *citA*, a 617 bp fragment was amplified using primers OMB173 and OMB174, flanked by an *Hin*dIII and a *Pst*I site. The second to amplify the downstream region of *citA*, a 567 bp fragment was amplified using primers OMB175 and OMB176, flanked by a *Pst*I site and an *Eco*RI site. These two fragments were first digested with appropriate restriction enzymes and then triple ligated into pNTPS138 (M.R.K. Alley, Imperial College London, unpublished) previously restricted with *Eco*RI*/Hin*dIII.

#### pMB288 (pNTPS138-Δ*citB*)

The plasmid construct used to delete *citB* (*CCNA_03757*) was made by PCR amplification of two fragments. The first to amplify the upstream region of *citB*, a 550 bp fragment was amplified using primers OMB184 and OMB185, flanked by a *Hin*dII and an *Nde*I. The second to amplify the downstream region of *citB*, a 538 bp fragment was amplified using primers OMB186 and OMB187, flanked by a *Nde*I site and an *Eco*RI site. These two fragments were first digested with appropriate restriction enzymes and then triple ligated into pNTPS138 (M.R.K. Alley, Imperial College London, unpublished) previously restricted with *Eco*RI*/Hin*dIII.

#### pMB289 (pNTPS138-Δ*citC*)

The plasmid construct used to delete *citC* (*CCNA_03758*) was made by PCR amplification of two fragments. The first to amplify the upstream region of *citC*, a 568 bp fragment was amplified using primers OMB188 and OMB189, flanked by a *HindII* and an *NdeI*. The second to amplify the downstream region of *citC*, a 551 bp fragment was amplified using primers OMB190 and OMB191, flanked by a *Nde*I site and a *Eco*RI site. These two fragments were first digested with appropriate restriction enzymes and then triple ligated into pNTPS138 (M.R.K. Alley, Imperial College London, unpublished) previously restricted with *Eco*RI*/Hin*dIII.

#### pMB309 (pNTPS138-Δ*citB*/*citC*)

The plasmid construct used to delete *citB* and *citC* (*CCNA_03757 and CCNA_03758*) was made by digestion of the upstream region of *citB* of the pMB288, a 532 bp fragment using the *Nde*I and *Eco*RI site. This fragment was ligated into the pMB289 digested by *Mfe*I and *Nde*I enzymes.

#### pMB302 (pMT335-*citA*)

The *citA* coding sequence was PCR amplified from NA1000 using the OMB179 and OMB182 primers. This fragment was digested with *Nde*I/*Eco*RI and cloned into *Nde*I/*Eco*RI-digested pMT335.

#### pMB303 (pMT335-*citB*)

The *citB* coding sequence was PCR amplified from NA1000 using the OMB194 and OMB195 primers. This fragment was digested with *Nde*I/*Eco*RI and cloned into *Nde*I/*Eco*RI-digested pMT335.

#### pMB304 (pMT335-*citC*)

The *citC* coding sequence was PCR amplified from NA1000 using the OMB196 and OMB197 primers. This fragment was digested with *Nde*I/*Eco*RI and cloned into *Nde*I/*Eco*RI-digested pMT335.

#### pMB310 (pMT335-*gltA*)

The *gltA* coding sequence was PCR amplified from *E. coli* MG1655 using the OMB203 and OMB204 primers. This fragment was digested with *Nde*I/*Eco*RI and cloned into *Nde*I/*Eco*RI-digested pMT335.

#### pMB287 (pSC-*citA*)

The *citA* coding sequence was PCR amplified from *C. crescentus* using the OMB179 and OMB183 primers. This fragment was digested with *Nde*I/*Hin*dIII and cloned into *Nde*I/*Hin*dIII digested pSC.

#### pMB325 (pMT335-*citA^H303W^*)

The *citA* catalytic mutant was generated using QuickChange Site-directed Mutagenesis kit (Agilent technologies). Briefly, the plasmid pMB302 (pMT335-*citA*) was PCR amplified using the mutagenic primers OMB232 and OMB233, containing the H303W mutation. This PCR was followed by a *Dpn*I digestion allowing to digest the parental plasmid and this digestion was used to transform electrocompetent *E. coli*. The integration of the site-directed mutation in *citA* coding sequence was verified by sequencing.

#### pMB326 (pMT335-*citA^H303A^*)

The *citA* catalytic mutant was generated using QuickChange Site-directed Mutagenesis kit (Agilent technologies). Briefly, the plasmid pMB302 (pMT335-*citA*) was PCR amplified using the mutagenic primers OMB236 and OMB237, containing the H303A mutation. This PCR was followed by a *Dpn*I digestion allowing to digest the parental plasmid and this digestion was used to transform electrocompetent *E. coli*. The integration of the site-directed mutation in *citA* coding sequence was verified by sequencing.

## AKNOWLEGMENTS

We thank Justine Collier, Sean Crosson, Martin Thanbichler, Michael Laub, Urs Jenal and Lucy Shapiro for materials, Julien Prados for help with Tn-Seq and suppressors analyses and Gaël Panis, Nicolas Kint for critical reading of the manuscript. This paper was supported by the Swiss National Science Foundatrion grant 31003A_182576 to Patrick H Viollier.

## AUTHOR CONTRIBUTION

MB, Conception and design, Acquisition of data, Analysis and interpretation of data, Drafting or revising the article, Contributed unpublished essential data or reagents. JP, VG Conception and design, Acquisition of data, Analysis and interpretation of data, Drafting or revising the article, Contributed unpublished essential data or reagents. LD, Conception and design, Acquisition of data, Analysis and interpretation of data, Contributed unpublished essential data or reagents. SR, Conception and design, Analysis and interpretation of data, Drafting or revising the article. PHV, Conception and design, Analysis and interpretation of data, Drafting or revising the article.

## SUPPLEMENTAL MATERIALS AND FIGURES

**Supplemental Table S3.**
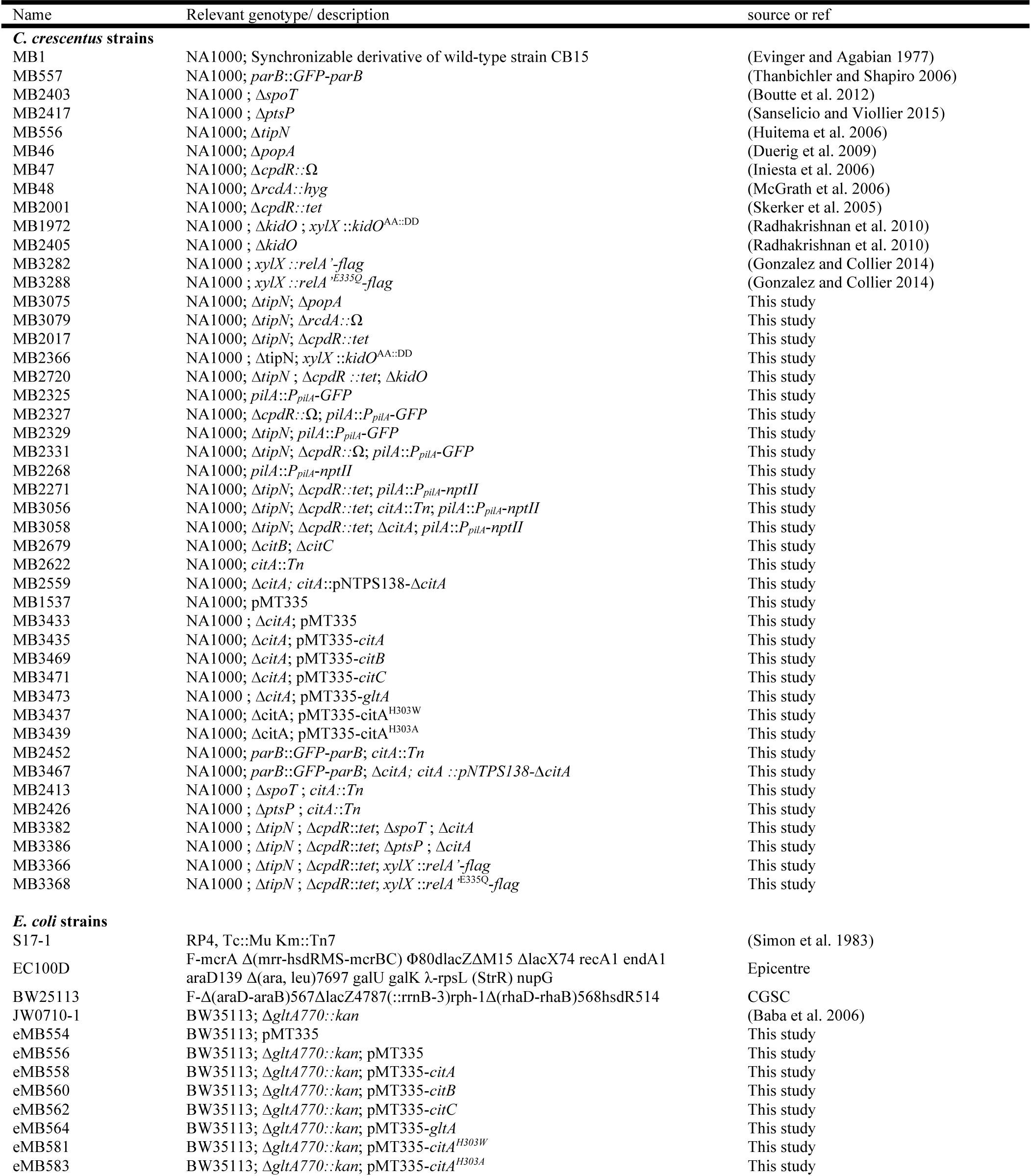
*C. crescentus* and *E. coli* strains

**Supplemental Table S4.**
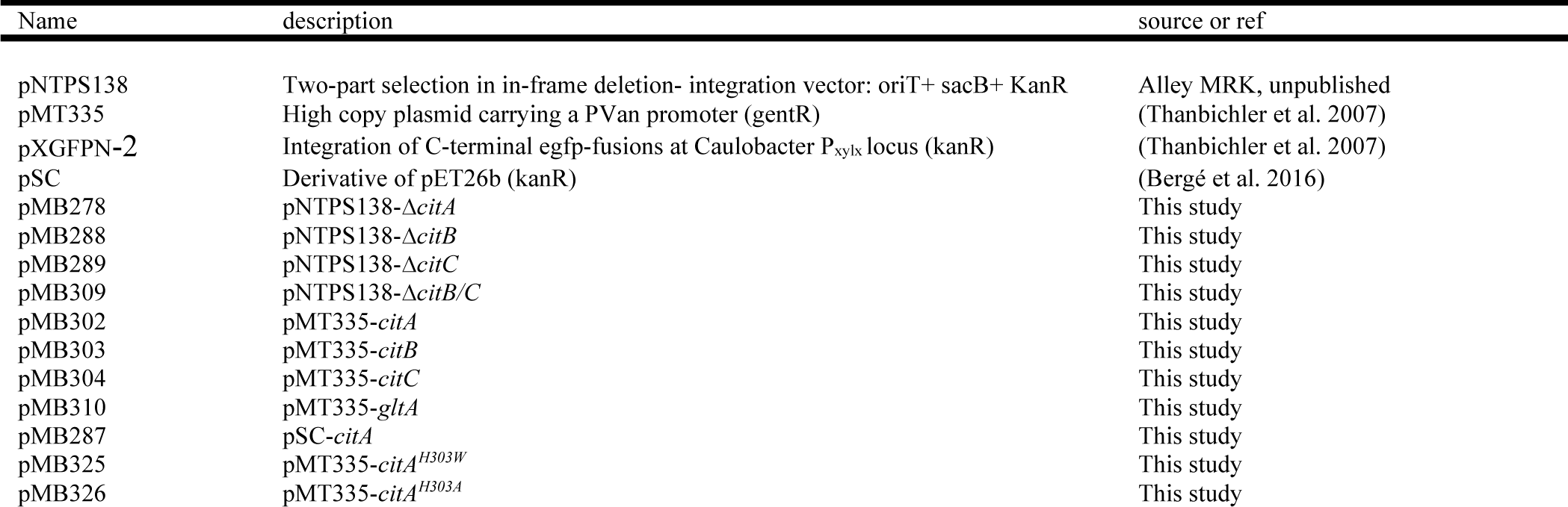
plasmids

**Supplemental Table S5.**
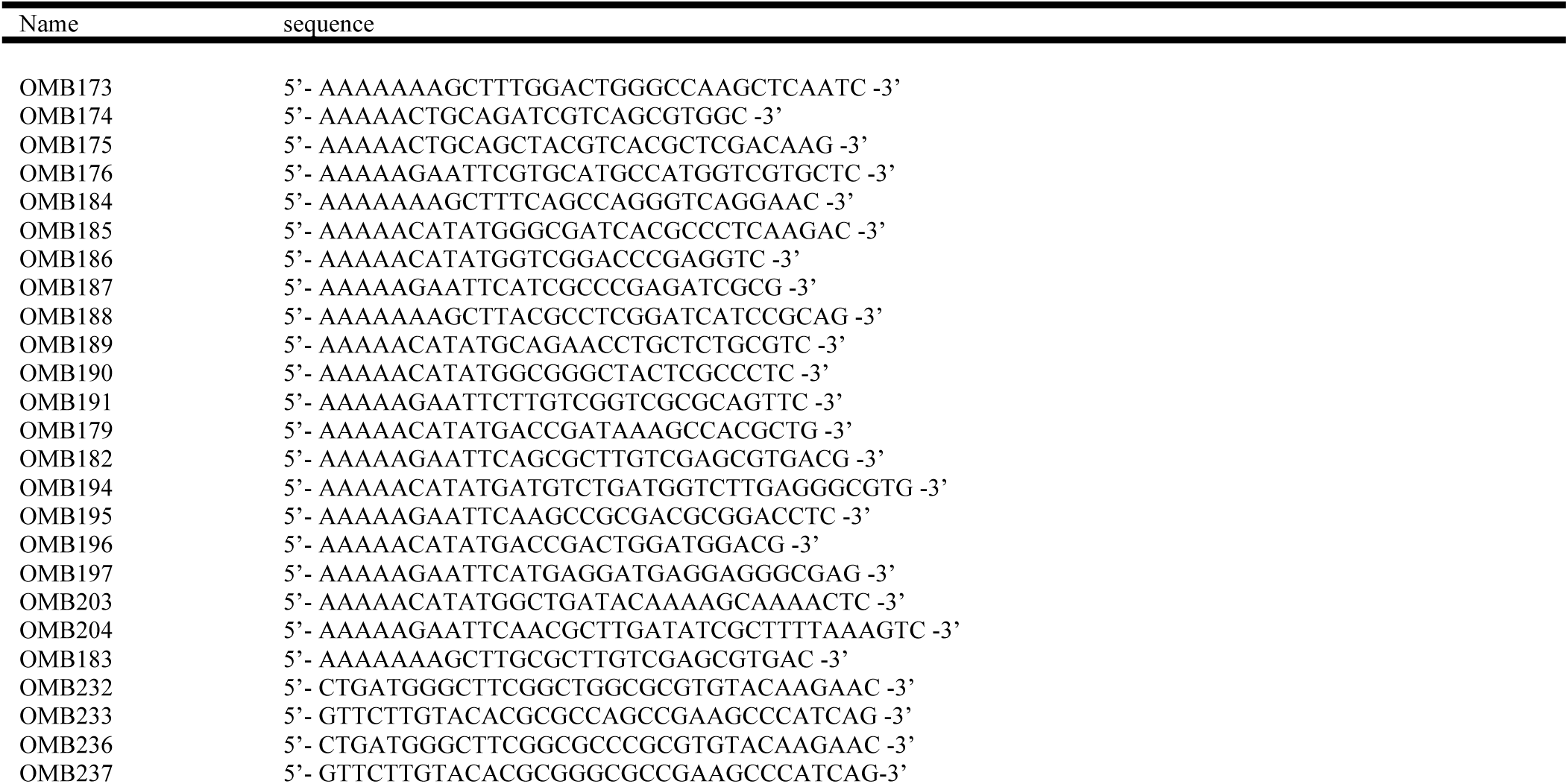
primers

**Figure 1- Figure supplemental 1.**
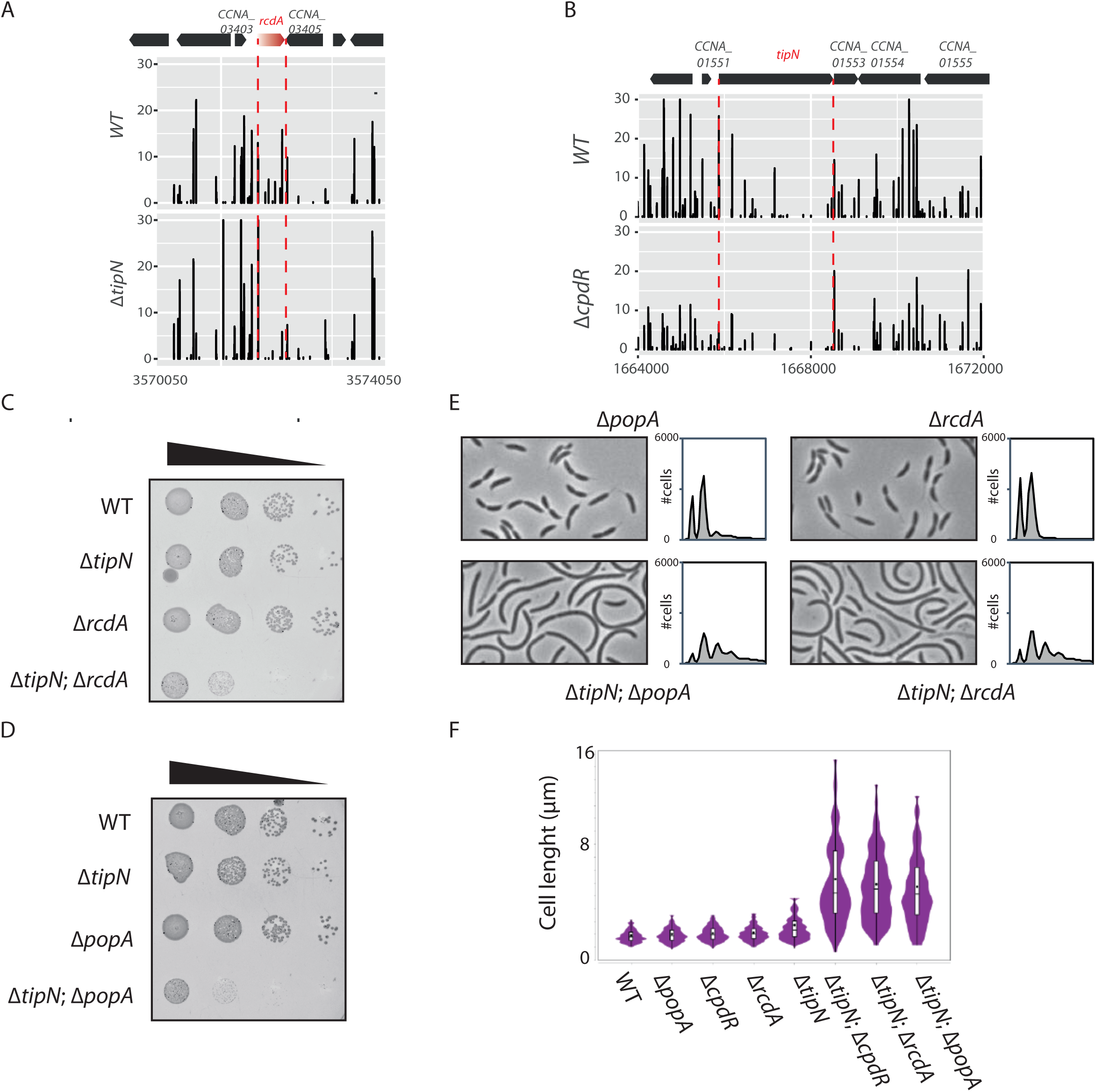
(A) Transposon libraries were generated in the *WT* and the Δ*tipN* mutant (MB556). The region of the *rcdA* locus is depicted. (B) Transposon libraries were generated in the *WT* and the Δ*cpdR* mutant (MB2001). The *tipN* coding sequence is represented showing a decrease in Tn insertions in the two mutants compared to the *WT*. (C) Spot dilutions of the indicated strains (MB1, MB556, MB48, MB3079 from top to bottom) done as described in Figure 1C. (D) Spot dilutions of the indicated strains (MB1, MB556, MB46, MB3075 from top to bottom) done as described in Figure 1 panel C. (E) Flow cytometry profiles and phase contrast images of *WT* (MB1), Δ*popA* (MB46), Δ*rcdA* (MB48), Δ*tipN* Δ*popA* (MB3075) or Δ*tipN* Δ*rcdA* (MB3079) double mutants. Genome content was analyzed by FACS during exponential growth in PYE. (F) Cell size distribution of *WT* (MB1), Δ*popA* (MB46), Δ*cpdR* (MB2001), Δ*rcdA* (MC48), Δ*tipN* (MB556), Δ*tipN* Δ*cpdR* (MB2017), Δ*tipN* Δ*rcdA* (MB3079) and Δ*tipN* Δ*popA* (MB3075). Strains were grown in PYE media. The cell length was measured automatically using MicrobeJ.

**Figure 1- Figure supplemental 2.**
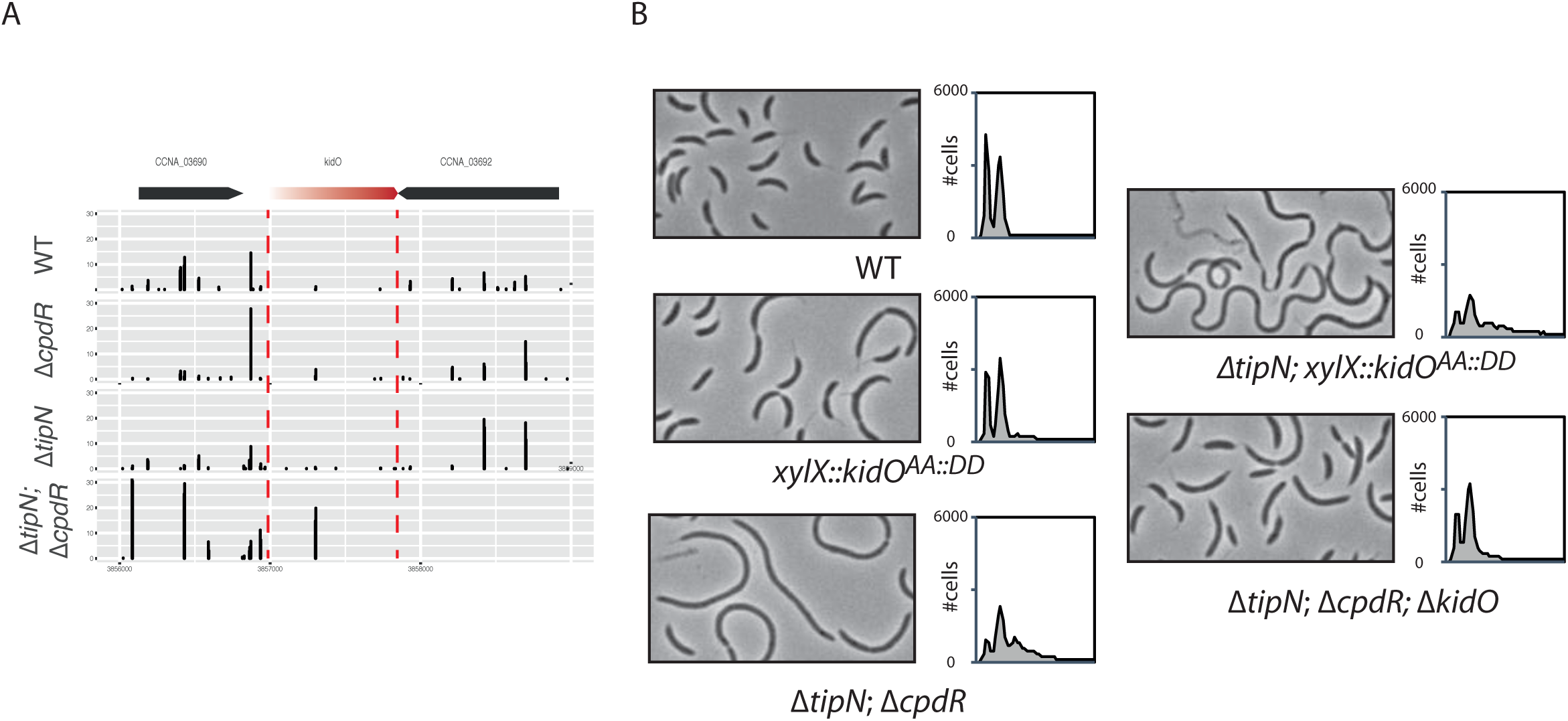
(A) Transposon libraries were generated in the *WT*, Δ*cpdR* (CC2001), Δ*tipN* (MBC556) single mutants or Δ*tipN* Δ*cpdR* double mutant (MB2017). The sites of Tn insertion were identified by deep sequencing and mapped onto the *C. crescentus* NA1000 reference genome. The *kidO* locus is depicted. The height of each line reflects the number of sequencing reads at this position. Tn insertions in *kidO* was increased in the Δ*tipN* Δ*cpdR* double mutant compared to the *WT* or the Δ*tipN* and Δ*cpdR* single mutant. (F) Flow cytometry profiles and phase contrast images of *WT* (MB1), Δ*tipN* Δ*cpdR* double mutant (MB2017), Δ*tipN* Δ*cpdR* Δ*kidO* triple mutant (MB2720) and the *WT* (MB1972) or Δ*tipN* (MB2366) expressing a non-degradable version of KidO (KidO^AA::DD^) under the control of the xylose promoter at the *xylX* locus. Genome content was analyzed by FACS during exponential growth in PYE. Note that the expression of KidO^AA::DD^ was not induced with xylose since the leakage of P_xyl_ was sufficient to induce strong filamentation in the double mutant Δ*tipN* Δ*cpdR*.

**Figure 2- Figure supplemental 1.**
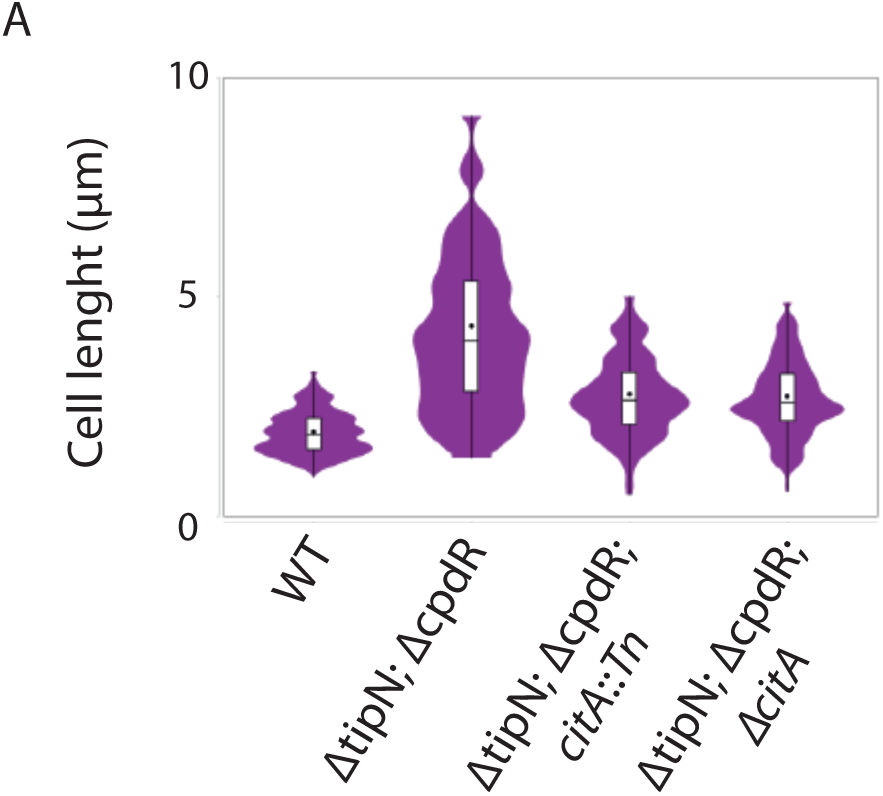
(A) Cell size distribution of *WT* (MB2268) (n=638), Δ*tipN* (MB2271) Δ*cpdR* double mutant (n=635), Δ*tipN* Δ*cpdR citA*::Tn triple mutant cells (MB3056) (n=553); and Δ*tipN* Δ*cpdR* Δ*citA* triple mutant cells (MB3058) (n=498). Strains were grown in PYE media. The cell length was measured automatically using MicrobeJ.

**Figure 3- Figure supplemental 1.**
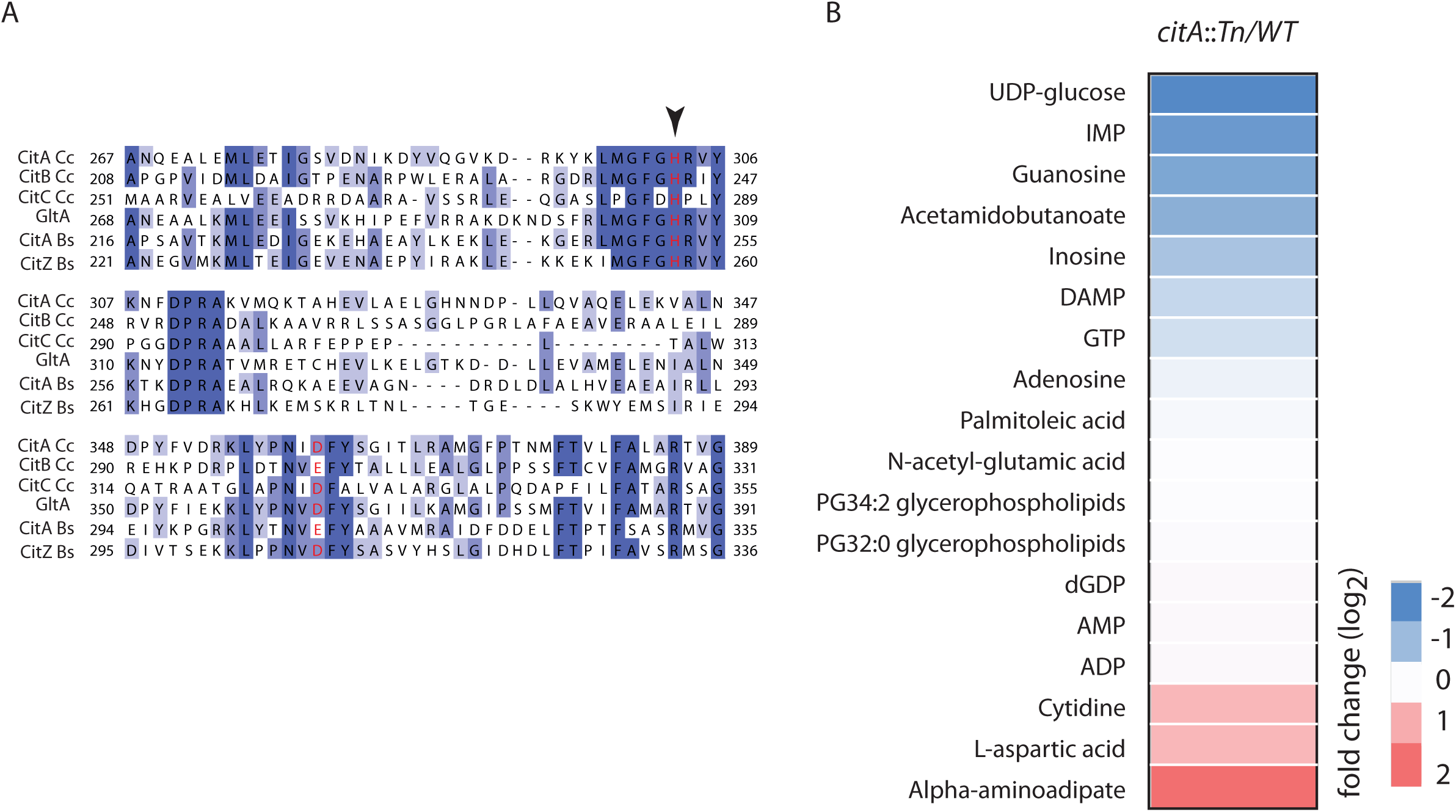
(A) Partial alignment of the active site of CitA (A0A0H3C985) with CitB (A0A0H3CCE2) and CitC (A0A0H3CD20) from *C*. *crescentus*, GltA (P0ABH7) from *E. coli*, CitA (P39119) and CitZ (P39120) from *Bacillus subtilis*. The histidine and aspartic acid catalytic site are highlighted in red. Arrow indicates the alanine or tryptophan substitution abolishing the catalytic activity of CitA (figure 4). (B) Heatmap showing the changes in the level of various metabolites among the *WT* and *citA*::Tn as measured by LC-MS. Cells were grown on PYE medium. Only the metabolites that were significantly increased or decreased (p-value<0.05) in Δ*citA* compared to *WT* cells are shown. Fold changes were calculated based on the mean of normalized ion counts from three biological replicates.

**Figure 4- Figure supplemental 1.**
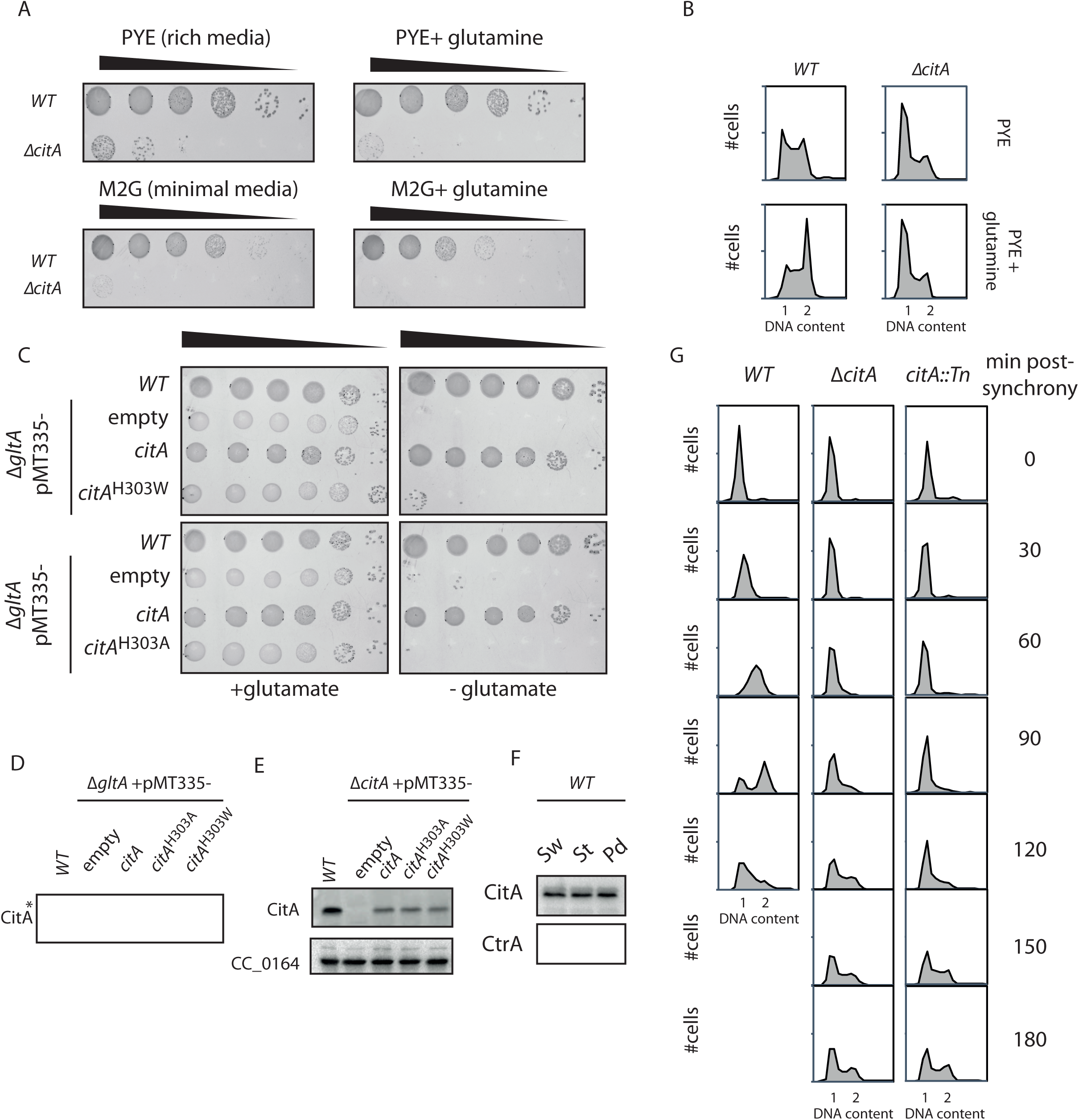
(A) Spot dilutions of the WT (MB1) and Δ*citA* (MB2559). The two strains were grown overnight in PYE, adjusted at an OD600 of 0.5 and serially diluted on PYE plate (upper part) or minimal medium M2G (lower part) containing (right panel) or not (left panel) glutamine. Eight microliters of each dilutions were spotted onto plates. (B) Flow cytometry profiles of WT (MB1) and Δ*citA* (MB2559). Genome content was analyzed by FACS during exponential phase in PYE (upper panel) or in PYE containing glutamine (lower panel). (C) Spot dilutions of the WT *E. coli* carrying an empty plasmid (eMB554) or *E. coli* Δ*gltA* cells harboring an empty plasmid (eMB556) or expressing *citA*^H303A^ (eMB583) or *citA*^H303W^ (eMB581) on minimal medium containing (left panel) or not (right panel) glutamate. Only the strain carrying a functional citrate synthase could growth without glutamate. (D) Immunoblot showing the abundance of CitA in the *E. coli* strains presented in panel C using antibody to CitA. Asterisk point to proteins that cross-react with the anti-CitA antibodies. All the CitA variants are expressed at similar levels. (E) Immunoblot showing the abundance of CitA in the *C. crescentus* strains presented in figure 4D using antibody to CitA. All the CitA variants are expressed at similar level. (F) Immunoblotting to determine the relative abundance of CitA and CtrA during the cell cycle of WT (MB1) *C. crescentus*. All strains were in synchronized in PYE and the Sw, St and Pd time point were taken at 0 min, 25 min and 60min respectively post-synchrony. (G) Flow cytometry profiles of the WT (MB1), *citA::Tn* (MB2622) and Δ*citA* (MB2559) to monitor DNA content throughout the *C. crescentus* cell cycle. WT (left panel), *citA::*Tn (middle panel) and Δ*citA* (right panel) were synchronized and samples were withdrawn every 30 minutes and prepared for FACS analysis.

**Figure 5- Figure supplemental 1.**
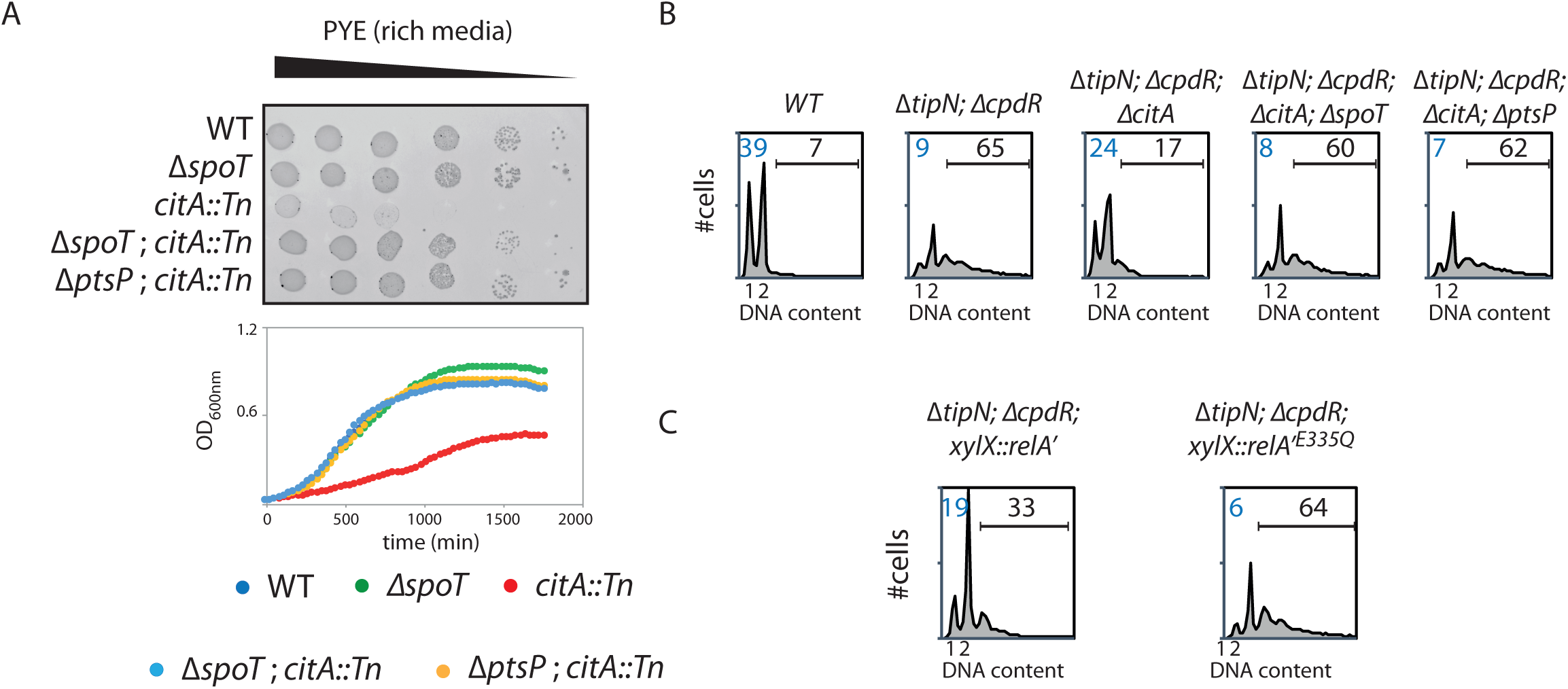
(A) Spot dilution and growth curve of the *WT* (MB1), Δ*spoT* (MB2403), *citA*::*Tn* (MB2622), Δ*spoT citA*::*Tn* (MB2413) and Δ*ptsP citA*::*Tn* (MB2426). For spot dilution, cells were grown overnight in PYE, adjusted to OD_600nm_ ∼0.5 and serially diluted on a rich PYE medium (upper part). For growth curve, cells were grown overnight in PYE, washed twice with M2 buffer and similar amount of each strain was used to inoculate PYE medium (left bottom part). (B) Flow cytometry profiles of *WT* (MB1), Δ*tipN* Δ*cpdR* (MB2017) double mutant, Δ*tipN* Δ*cpdR* Δ*citA* (MB3058) triple mutant, Δ*tipN* Δ*cpdR* Δ*citA* Δ*spoT* (MB3382) or Δ*tipN* Δ*cpdR* Δ*citA* Δ*ptsP* (MB3386) quadruple mutant. Genome content was analyzed by FACS during exponential phase in PYE. (C) Flow cytometry profiles of Δ*tipN*; Δ*cpdR* expressing RelA’ (MB3366) or RelA’^E335Q^ (MB3368) under the control of the xylose promoter at the *xylX* locus. Genome content was analyzed by FACS during exponential growth in PYE containing xylose for 5h to induce RelA’ or RelA’^E335Q^.

